# Harmonizing amyloid PET tracers at the local level: evaluation of regional Centiloid equations

**DOI:** 10.64898/2026.11.10.765694

**Authors:** Erica Silvestri, Pavanjit Chaggar, Susanna Gobbi, Roxana Aldea, Gregory Klein, Matteo Tonietto

## Abstract

**INTRODUCTION:** The Centiloid (CL) framework standardizes global amyloid-PET across radiotracers. While global scaling remains essential for participant selection, amyloid-targeting therapies require quantifying localized plaque removal. However, applying global equations regionally introduces systematic bias, particularly in subcortical structures where off-target binding alters non-specific background.

**METHODS:** We extended the Centiloid framework regionally and voxel-wise using GAAIN datasets: Level-1 ([^11^C]PiB) and Level-2 paired [^11^C]PiB and fluorinated head-to-head cohorts ([^18^F]florbetaben, [^18^F]florbetapir, [^18^F]flutemetamol, and [^18^F]NAV4694). Level-1 defines the regional Centiloid scale using region-specific intercepts while fixing the global slope. Level-2 calibrates fluorinated tracer uptake to reference [¹¹C]PiB SUVR using linear equations. We compared three slope-intercept models (Global, Regional-Intercept, and Regional) using information criteria and Mean Absolute Error (MAE).

**RESULTS:** Regional-Intercept model was identified as optimal across all four radiotracers. In neocortical regions, Global and Regional-Intercept models showed high agreement across canonical lobes. In deep gray matter, global equations introduced large systematic offsets, with mean absolute conversion errors in the pallidum reaching 27.7 to 45.6 CL. The Regional-Intercept model eliminated this bias, reducing subcortical MAE on average below 15 CL while preserving a unified biological scale.

**DISCUSSION:** Applying global Centiloid equations directly to regions introduces substantial bias in subcortical regions. With the Regional-Intercept model, the same Centiloid value corresponds to the same amyloid concentration across brain regions and radiotracers. This provides a reliable basis for cross-sectional regional evaluation, multi-tracer cohort pooling, tracking regional clearance, and mapping regional progression in clinical research.

## 1. Introduction

Positron emission tomography (PET) imaging of amyloid-β (Aβ) plaques is a cornerstone in the research, diagnosis, and management of Alzheimer’s disease (AD). Following the introduction of [^11^C]PiB, various [^18^F]-labelled radiotracers were developed and approved for clinical practice (Iaccarino *et al*., 2025). As the number of clinically available options expanded, it became essential to standardize quantitative PET measures across different imaging centers, radiotracers, and analysis pipelines.

To address this challenge, the Centiloid (CL) framework was established, providing a standardized, tracer- and pipeline-independent scale anchored to the average values in young healthy controls (YC, 0 CL) and typical mild-to-moderate AD patients (100 CL) (Klunk *et al*., 2015). In the standard CL approach, tracer-specific Standardized Uptake Value Ratios (SUVRs) are extracted from a predefined global cortical target region and converted into a single Centiloid value with a linear equation.

While this global framework remains the benchmark for overall plaque burden, amyloid-lowering therapies highlight the need for regional assessments. Emerging trials demonstrate regional plaque clearance differences among individuals treated with aducanumab (Chiao *et al*., 2019), gantenerumab (McCullough *et al*., 2025), or lecanemab (Iizuka, Watanabe and Kameyama, 2026; Noh *et al*., 2026). These studies rely on raw SUVRs obtained with different tracers and pipelines, making their results difficult to compare quantitatively. Postmortem evaluations further reveal patchy clearance patterns, where cleared structures sit juxtaposed against regions with limited plaque reduction (Boon *et al*., 2025; Brown *et al*., 2026). Regional measures are equally valuable in preclinical disease, where early focal deposition predicts tau spread and cognitive decline prior to global positivity (Farrell *et al*., 2018, 2024). A standardized regional framework would allow mapping how clearance progresses in each structure, to better understand how therapies work. Finally, linking regional plaque reduction to downstream tau spread and neurodegeneration can help explain why clinical benefit is greater in individuals with low initial tau burden, and how clearing specific regions relates to specific cognitive outcomes.

Despite the utility of regional assessments, applying global Centiloid equations directly to individual brain structures has received limited systematic evaluation. The standard conversion uses linear equations from the original two-level Centiloid protocol. These equations were calibrated for a global cortical composite volume of interest (Klunk *et al*., 2015), and not for individual brain regions. At the regional level, using the global equations raises methodological problems at both calibration levels. At Level-1, non-specific tracer uptake in young controls differs on average across brain areas. As a result, 0 CL might not correspond to zero amyloid in every region. At Level-2, each non-PiB tracer is converted with a single equation derived from the cortical target region. This ignores regional differences in tracer kinetics, off-target binding and tissue composition which could introduce systematic bias in regional values. This is particularly concerning in subcortical areas, where tracer-specific off-target binding differs between tracers and can change the linear relationship between the fluorinated tracer and [^11^C]PiB SUVRs.

In this study, we extend the Centiloid framework to the regional level at both calibration levels. Our goal is a regional scale on which the same Centiloid value corresponds to the same amyloid concentration, in any brain region and with any tracer. Like the global Centiloid scale, this regional framework is intended for measuring regional amyloid burden cross-sectionally or during disease progression or treatment, pooling multi-tracer cohorts in multicenter studies. At Level-1, we use healthy young controls to set the non-specific background signal of each region to zero, so that 0 CL reflects the absence of amyloids in all structures. At Level-2, we use head-to-head Global Alzheimer’s Association Interactive Network (GAAIN, (Klunk *et al*., 2015)) datasets for [^18^F]florbetaben, [^18^F]florbetapir, [^18^F]flutemetamol, and [^18^F]NAV4694 to compare three linear models: a Global model, which applies the standard composite equations to all structures; a Regional-Intercept model, which keeps the global slope but adds a region-specific intercept; and a fully Regional model, which fits a separate equation for each region. This comparison identifies the best approach for regional Centiloid harmonization.

## 2. Methods

### 2.1 Cohorts Description

Data were obtained from standard reference datasets provided by GAAIN (www.gaain.org/centiloid-project), comprising dynamic amyloid-PET and high-resolution T1-weighted (T1w) MRI. These cohorts fulfill the two calibration levels established by the Centiloid framework:

- Level-1 Calibration Dataset ([^11^C]PiB Anchoring, (Klunk *et al*., 2015)): Consisted of young controls (YC, ≤45 years; N=34) defining the 0 CL biological zero-anchor and clinically diagnosed AD patients (N= 45) representing the 100 CL high-amyloid anchor point.
- Level-2 Head-to-Head (H2H) Datasets: Included four paired cross-sectional cohorts for four fluorinated radiotracers: [^18^F]florbetaben (FBB, (Rowe *et al*., 2017)), [^18^F]florbetapir (FBP, (Navitsky *et al*., 2018)), [^18^F]flutemetamol (FMM, (Vandenberghe *et al*., 2010; Lowe *et al*., 2017)), and [^18^F]NAV4694 (NAV, (Rowe *et al*., 2016)). Each H2H dataset contains a YC group (N= 10 to 24 per tracer) alongside an elderly cohort (N= 25 to 50 per tracer) spanning the diagnostic spectrum (AD, Mild Cognitive Impairment (MCI), Frontotemporal Dementia (FTD), and Healthy Controls (HC)). Participants underwent paired scans using the fluorinated tracer and reference [^11^C]PiB (**Table 1**).

**Table 1.** Demographic characteristics and PET imaging acquisition protocols of the included datasets. The cohorts are structured according to the two primary validation tiers of the GAAIN Centiloid standardization framework. Level-1 Anchoring establishes the absolute regional zero-anchor bounds using standard [^11^C]PiB imaging in a cohort of amyloid-negative Young Controls (YC) and clinically confirmed Alzheimer’s Disease (AD) patients. Level-2 Head-to-Head (H2H) Harmonization datasets encompass cohorts of elderly participants across the clinical spectrum (including Mild Cognitive Impairment (MCI), Frontotemporal Dementia (FTD), and Healthy Controls (HC)) who received paired, closely-timed scans with both a fluorinated tracer and the reference [^11^C]PiB tracer. Corresponding scanning windows (minutes post-injection) and temporal frame configurations are detailed

| Tracer | Dataset Group | Elderly (n)<br>(AD / MCI / FTD / HC) | Young Controls (YC) | Acquisition Protocol |  | PET Frames |
|---|---|---|---|---|---|---|
| [<sup>11</sup>C]PiB | Level-1 Anchoring | 45 / 0 / 0 / 0 | 34 | [<sup>11</sup>C]PiB | 50–70 min | 4 x 5 min |
| [<sup>18</sup>F]Florbetaben | Level-2 H2H | 8 / 9 / 2 / 6 | 10 | [<sup>18</sup>F]FBB | 90–110 min | 4 x 5 min |
|  |  |  |  | [<sup>11</sup>C]PiB | 50–70 min | 4 x 5 min |
| [<sup>18</sup>F]Florbetapir | Level-2 H2H | 17 / 7 / 0 / 9 | 13 | [<sup>18</sup>F]FBP | 50–60 min | 2 x 5 min |
|  |  |  |  | [<sup>11</sup>C]PiB | 50–70 min | 4 x 5 min |
| [<sup>18</sup>F]Flutemetamol | Level-2 H2H | 20 / 20 / 0 / 10 | 24 | [<sup>18</sup>F]FMM | 90–110 min | 4 x 5 min |
|  |  |  |  | [<sup>11</sup>C]PiB | 50–70 min | 4 x 5 min |
| [<sup>18</sup>F]NAV4694 | Level-2 H2H | 7 / 10 / 3 / 25 | 10 | [<sup>18</sup>F]NAV | 50–70 min | 4 x 5 min |
|  |  |  |  | [<sup>11</sup>C]PiB | 50–70 min | 4 x 5 min |

### 2.2 Image Processing, Region Extraction, and Quality Control

Following the native space PET processing pipeline (Royse *et al*., 2021), SUVRs normalized to the whole cerebellum were extracted for the global cortical composite, five cortical regions (cingulate, frontal, occipital, parietal, and temporal), and five subcortical structures (caudate, hippocampus, pallidum, putamen, and thalamus). A rigorous visual quality control across all preprocessing, registration, and spatial normalization steps was performed prior to calculating Centiloid values. If misalignment or registration artifacts were detected, parameter fine tuning or targeted re-registration corrections were executed to achieve optimal alignment. T1w technical exclusion criteria comprise severe spatial anisotropy (non-isotropic slice thickness ≥ 5.0mm), contrast degradation precluding gray/white matter segmentation, or gross parcellation failures. Full image processing and spatial normalization details provided in **Supplementary Materials**.

### 2.3 Global Centiloid Calibration and Pipeline Validation

Global scaling equations were derived following GAAIN guidelines (Klunk *et al*., 2015):

- Level-1 Pipeline Validation: Global CL_PiB_ values from our pipeline were regressed against published GAAIN benchmarks using the Level-1 [^11^C]PiB anchoring dataset, meeting standard acceptance criteria (R^2^≥0.98, slope 1.00±0.02, intercept 0.00±2.00 CL).
- Level-2 Head-to-Head Calibration: Paired global cortical SUVRs ([^11^C]PiB vs. [^18^F]tracer) were regressed across all four H2H datasets to derive global conversion parameters, meeting the GAAIN threshold of R^2^≥0.70.

Through global validation, the derived slopes and intercepts were used to establish regional and voxel-wise frameworks.

### 2.4 Regional Centiloid Framework

Our regional framework aims to ensure that a given Centiloid value reflects the same amyloid burden in every brain region, regardless of tracer, scanner or pipeline. This requires two steps: anchoring 0 CL to the non-specific background signal of each region at Level-1, and making each tracer equivalent to reference [¹¹C]PiB SUVR at Level-2.

#### 2.4.1 Level-1: Centiloid Scale Definition

In healthy young controls, reference [¹¹C]PiB SUVR values vary across brain regions due to differences in non-displaceable signal rather than amyloid deposition (**Figure 1A**). To ensure that 0 CL represents the absence of amyloid in all structures, we anchored each region to its own young control average using a region-specific intercept (*intercept_CL_*_,*r*_). For the 100 CL anchor, we did not scale each region separately. Brain structures differ in the maximum amount of amyloid they can accumulate (their amyloid carrying capacity) (Whittington, Sharp and Gunn, 2018). Scaling each region separately between 0 and 100 would stretch the scale differently in each region and distort regional comparisons. Instead, we kept a single global slope (*slope_CL_*_,*g*_), derived from the standard global target composite, for all regions. With a shared slope, a region that can hold less amyloids simply reaches lower Centiloid values, instead of being artificially stretched to the same maximum. In this way, a given Centiloid value reflects the same amyloid concentration in all regions.

**Figure 1.**
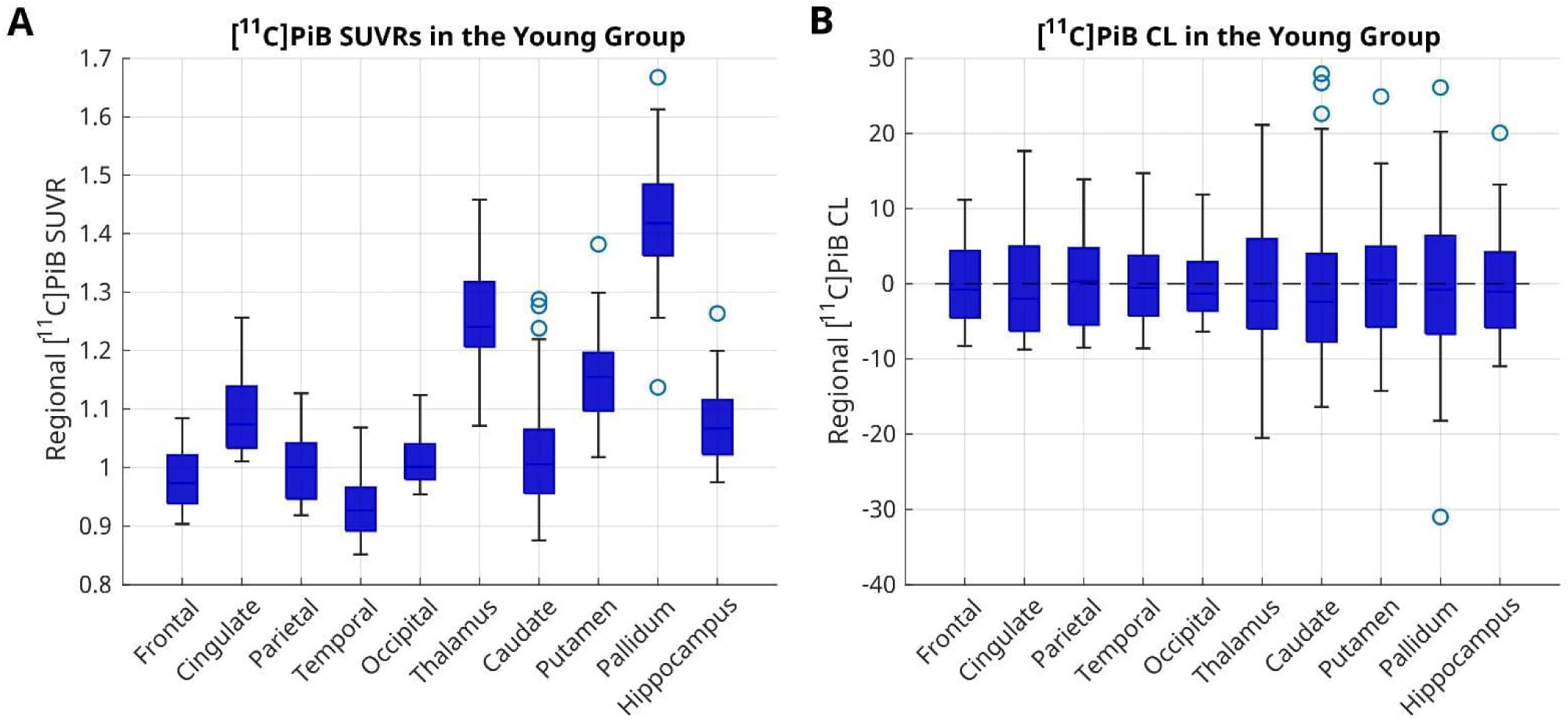
Regional zero-anchor (0 CL) definition from the young controls group. Panel A, boxplots of regional [^11^C]PiB standard uptake value ratios (SUVR) in the young control group of the anchoring dataset across ten cortical lobes and subcortical deep gray matter regions of interest. Panel B, regional Centiloid (CL) values distributions in the same young control cohort following the introduction of the region-specific intercept at Level-1, successfully centering the young control group near 0 CL across both cortical and subcortical regions. Boxes denote the interquartile range (IQR), central horizontal lines represent the median value, whiskers extend to 1.5xIQR, and open circles indicate individual outliers.

Mathematically, the uniform global slope (*slope_CL_*_,*g*_) was calculated from the standard global cortical composite:

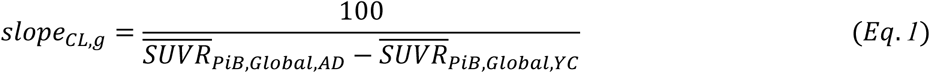

where the 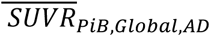 and 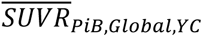 represent the mean global target region [^11^C]PiB SUVR of the AD and YC cohorts from the Level-1 calibration dataset.

The regional Centiloid value (*CL_r_*) for region *r* is computed as:

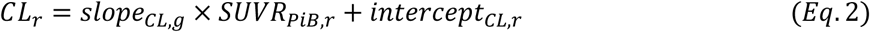

where the region-specific intercept (*intercept_CL_*_,*r*_) anchors the mean [^11^C]PiB SUVR of the Level-1 YC cohort in region 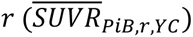 to 0 CL:

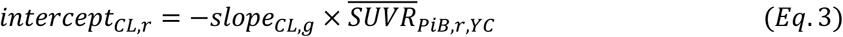

#### 2.4.2 Level-2: Regional Head-to-Head Calibration Models

Level-2 harmonization must resolve differences in tracer-specific properties, such as off-target binding and dynamic range, which vary across brain structures. Level-2 calibration maps regional fluorinated tracer uptake (*SUVR*_18*F*,*r*_) to the reference [^11^C]PiB scale (*SUVR*_11*C*,*r*_), ensuring that regional Centiloid values represent the same amyloid burden across tracers. Following GAAIN guidelines, linear regressions were fitted with *SUVR*_18*F*,*r*_ as the dependent variable and *SUVR_PiB_*_,*r*_ as the independent variable using unweighted linear least squares, pooling young controls and elderly cohorts for each head-to-head dataset. We evaluated three competing model formulations for regional mapping:

1. **Global Equation:** Assumes spatial homogeneity across regions and applies the single global slope (*slope_G_*) and global intercept (*intercept_G_*) derived from global calibration (from the standard global target composite mask) to all individual ROIs:

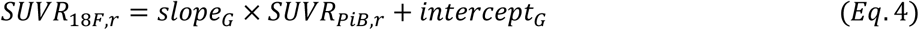
2. **Regional-Intercept Equation:** Assumes that the slope is the same across regions, but the intercept can vary due to different off-target binding and other technical factors. This hybrid approach keeps the unified global slope (*slope_G_*), but introduces region-specific intercepts (*intercept_G_*_,*r*_) calculated from local ROI data to correct local offsets:

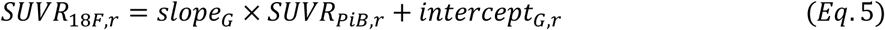
3. **Regional Equation:** Fits independent slopes (*slope_r_*) and intercept (*intercept_r_*) for each specific region using only that region’s data:

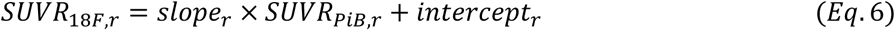

#### 2.4.3 Level-2: Model Selection

To determine the optimal mathematical approach for region-wise cross-tracer calibration, model performance was evaluated in terms of the balance between mathematical goodness-of-fit, parsimony, and error reduction directly on the CL values across the head-to-head datasets.

Following GAAIN guidelines, each calibration regression (Eqs. 4 to 6) used the fluorinated SUVR as the dependent variable and the [^11^C]PiB SUVR as the independent variable. The model fit residuals of these regressions therefore measure errors in fluorinated SUVR units rather than in [^11^C]PiB SUVR or Centiloid space. To evaluate candidate models on an identical, clinically interpretable scale, each calibration equation was inverted to compute [^11^C]PiB-equivalent SUVRs. These equivalent values and measured *SUVR_PiB_*_,*r*_ were then converted into regional Centiloid units *CL_PiB_*_,*r*_ and predicted 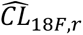 using the Level-1 equations. Model performance was then quantified using the following metrics:

- **Information Criteria:** To evaluate the trade-off between predictive accuracy and parsimony, Akaike Information Criterion (AIC) and Bayesian Information Criterion (BIC) were computed from residual sum of squares calculated directly on the Centiloid scale across pooled observations (participants (*P*) x regions (*R*)). Under the assumption of normally distributed residuals with constant variance, AIC and BIC were calculated as:

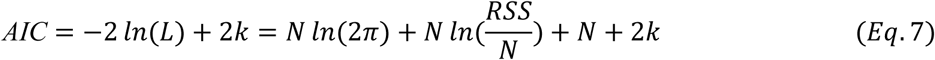

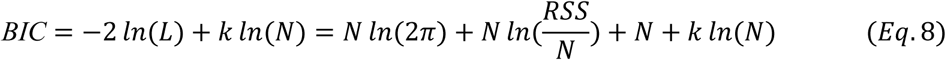

where *L* is the maximized value of the likelihood function, *k* is the number of estimated parameters including the residual variance, and *N* is the total number of pooled observations (*N = P x R*). The residual sum of squares (*RSS*) was calculated directly on the Centiloid scale:

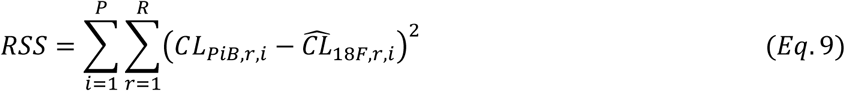

- **Mean Absolute Error (MAE):** Quantified conversion error in Centiloid units at the regional level (*MAE_H_*_2*H*,*r*_) for each ROIs and at the overall head-to-head dataset level (*MAE_H2H_*):

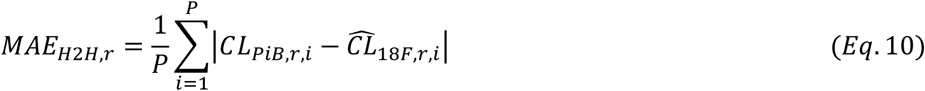

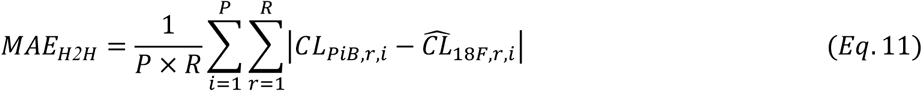

Regional and H2H-level MAE profiles across all three models were benchmarked against the native global cortical composite MAE to compare regional vs global variability.

Quantitative model selection relied on AIC as the primary criterion, supported by BIC and dataset-level MAE. To evaluate spatial goodness of fit alongside these numerical metrics, cross-tracer scatterplots comparing regional fluorinated against [^11^C]PiB SUVRs were visually inspected across cortical lobes and subcortical nuclei alongside fitted regression lines. These visual evaluations served as qualitative diagnostic checks and were not used to make model selection decisions.

Statistical analyses and model evaluations were performed in MATLAB (version R2022b, MathWorks).

## 3. Results

### 3.1 Characteristics of the Cohorts and Technical Exclusions

Demographic characteristics, clinical diagnoses, tracer-specific acquisition protocols, and sample sizes for all included datasets are summarized in **Table 1** and detailed in **Table S1**. Following automated processing and visual quality control, four scans were excluded due to FreeSurfer segmentation failures:

- Level-1 [^11^C]PiB Anchoring Dataset: Two AD participants were excluded from regional parcellation analyses due to structural MRI degradation. Their global target SUVR values were imputed to preserve the standard GAAIN reference sample size for global equation recalculation (Supplementary Methods).
- Level-2 [^18^F]FBP Dataset: Two AD participants were excluded: one due to thick sagittal slicing (5.2mm) violating FreeSurfer isotropic resolution requirements, and the other due to severe gray-to-white matter contrast degradation.

All remaining datasets passed quality control and were included in the regional and voxel-wise harmonization analyses.

### 3.2 Global Centiloid Calibration and Pipeline Validation

Technical validation of our dedicated processing pipeline was performed using the standard GAAIN [^11^C]PiB anchoring dataset. Global Centiloid values derived from our pipeline showed high agreement with the published GAAIN reference values anchoring dataset (**Figure S1**, R^2^=0.99, slope 1.01, intercept=-0.33 CL), meeting standard Level-1 acceptance criteria. For the Level-2 global calibrations, linear regressions comparing global cortical SUVRs from individual scans consistently exceeded the validation threshold (R^2^≥0.70 across all four head-to-head fluorinated tracer datasets, **Figure S2**).

### 3.3 Regional Centiloid Framework

#### 3.3.1 Level-1 Calibration: Definition of Regional Anchor Point 0

In the YC cohort, raw [^11^C]PiB SUVRs exhibited significant regional variance (**Figure 1A**), with mean values ranging from 0.93 SUVR in the temporal lobe to 1.15-1.43 SUVR in deep gray matter structures (putamen, thalamus, and pallidum). Following Level-1 calibration, regional values centered at 0 CL across all ten evaluated regions (**Figure 1B**). This adjustment aligned the regional non-specific uptake to a uniform zero floor across both cortical and subcortical structures.

#### 3.3.2 Level-2: Regional Head-to-Head Harmonization

Within cortical regions, SUVRs showed strong linear correlation across all evaluated tracers (R^2^ range: 0.86−0.98; **Figure S3**). In these cortical regions, [^18^F]NAV4694 exhibited the highest correlation with [^11^C]PiB (R^2^=0.93−0.98), followed by [^18^F]FMM (R^2^=0.94−0.96), [^18^F]FBB (R^2^=0.93−0.96), and [^18^F]FBP (R^2^=0.86−0.91). In contrast, subcortical structures exhibited weaker correlations (R^2^ range: 0.24−0.93). While correlations remained high in structures such as the putamen (R^2^=0.82−0.93), the match with [^11^C]PiB SUVR degraded in subcortical structures. This decrease was particularly pronounced in the pallidum, where correlation coefficients fell to R^2^=0.24 for [^18^F]FBP and R^2^=0.30 for [^18^F]FBB.

Using AIC as the main criterion, supported by BIC and MAE, the Regional-Intercept model was identified as the optimal framework across all four radiotracers (**Table 2**). AIC favored the Regional-Intercept model in every dataset, providing the best balance between accuracy and simplicity. BIC supported this selection for [^18^F]FBB, [^18^F]FBP, and [^18^F]FMM, but favored the simpler Global model for [^18^F]NAV. This reflects the high binding concordance and minimal non-specific background of [^18^F]NAV, which make its regional values less sensitive to intercept adjustments.

**Table 2.** Statistical model selection results for regional Centiloid standardization across fluorinated tracers. Goodness-of-fit and parsimony metrics are compared across three linear frameworks evaluated on Level-2 head-to-head datasets: (1) a *Global* model applying uniform scaling across all structures, (2) the *Regional-Intercept* model incorporating a shared global slope alongside region-specific intercepts, and (3) a *Regional* model mapping unique slopes and intercepts for every structure. N denotes the total number of samples (calculated as the number of included participants multiplied by the number of regions of interest). AIC, Akaike Information Criterion; BIC, Bayesian Information Criterion; MAE_H2H_ Mean Absolute Error computed in Centiloid units. The optimal model for each tracer, determined by the lowest AIC, is indicated in bold.

| Tracer | Model | Slope | Intercept | Parameters | N | AIC | BIC | MAE<sub>H2H</sub> |
|---|---|---|---|---|---|---|---|---|
| [<sup>18</sup>F]Florbetaben | Global | Global | Global | 2 | 350 | 2944.1 | 2951.8 | 11.0 |
|  | ***Regional-Intercept*** | ***Global*** | ***Regional*** | ***11*** | 350 | ***2817.7*** | ***2860.1*** | ***9.4*** |
|  | Regional | Regional | Regional | 20 | 350 | 2957.6 | 3034.7 | 10.5 |
| [<sup>18</sup>F]Florbetapir | Global | Global | Global | 2 | 440 | 4104.7 | 4112.9 | 19.3 |
|  | ***Regional-Intercept*** | ***Global*** | ***Regional*** | ***11*** | 440 | ***3980.4*** | ***4025.4*** | ***16.2*** |
|  | Regional | Regional | Regional | 20 | 440 | 4108.7 | 4190.4 | 17.5 |
| [<sup>18</sup>F]Flutemetamol | Global | Global | Global | 2 | 740 | 6590.0 | 6599.2 | 13.7 |
|  | ***Regional-Intercept*** | ***Global*** | ***Regional*** | ***11*** | 740 | ***5635.6*** | ***5686.3*** | ***6.5*** |
|  | Regional | Regional | Regional | 20 | 740 | 5670.0 | 5762.1 | 6.5 |
| [<sup>18</sup>F]NAV4694 | Global | Global | Global | 2 | 550 | 4068.5 | 4077.1 | 5.8 |
|  | ***Regional-Intercept*** | ***Global*** | ***Regional*** | ***11*** | 550 | ***4042.5*** | ***4089.9*** | ***5.3*** |
|  | Regional | Regional | Regional | 20 | 550 | 4130.1 | 4216.3 | 5.6 |

Regional model performance was evaluated by comparing linear fits against empirical scatterplot data expressed in native SUVRs across neocortical lobes (**Figure 2**) and subcortical structures (**Figure 3**). In neocortical regions, regression fits for the Global and Regional-Intercept models showed high concordance. Specifically, the frontal lobe demonstrated statistical equivalence between the two models across all four tracers (inter-model differences within 1.3 CL; TOST equivalence test with a ±2.5 CL margin, all p < 0.001). Parietal differences remained modest across cohorts (−0.4 to 2.8 CL), satisfying equivalence for three of four tracers (p < 0.001 for [^18^F]FBP, [^18^F]FMM, and [^18^F]NAV, while exceeding the margin for [^18^F]FBB at 2.8 CL). In contrast, cingulate and temporal lobes exhibited tracer-dependent divergence, exceeding the equivalence boundary with offsets up to 4.7 CL for [^18^F]FMM in the cingulate and −6.95 CL for [^18^F]FBP in the temporal lobe. In the occipital lobe (which is excluded from the standard GAAIN composite definition) inter-model differences between the Global and Regional-Intercept models ranged from −1.4 to 6.4 CL, while overall differences in conversion accuracy (MAE) between the two models remained under 1.1 CL across all tracers.

**Figure 2.**
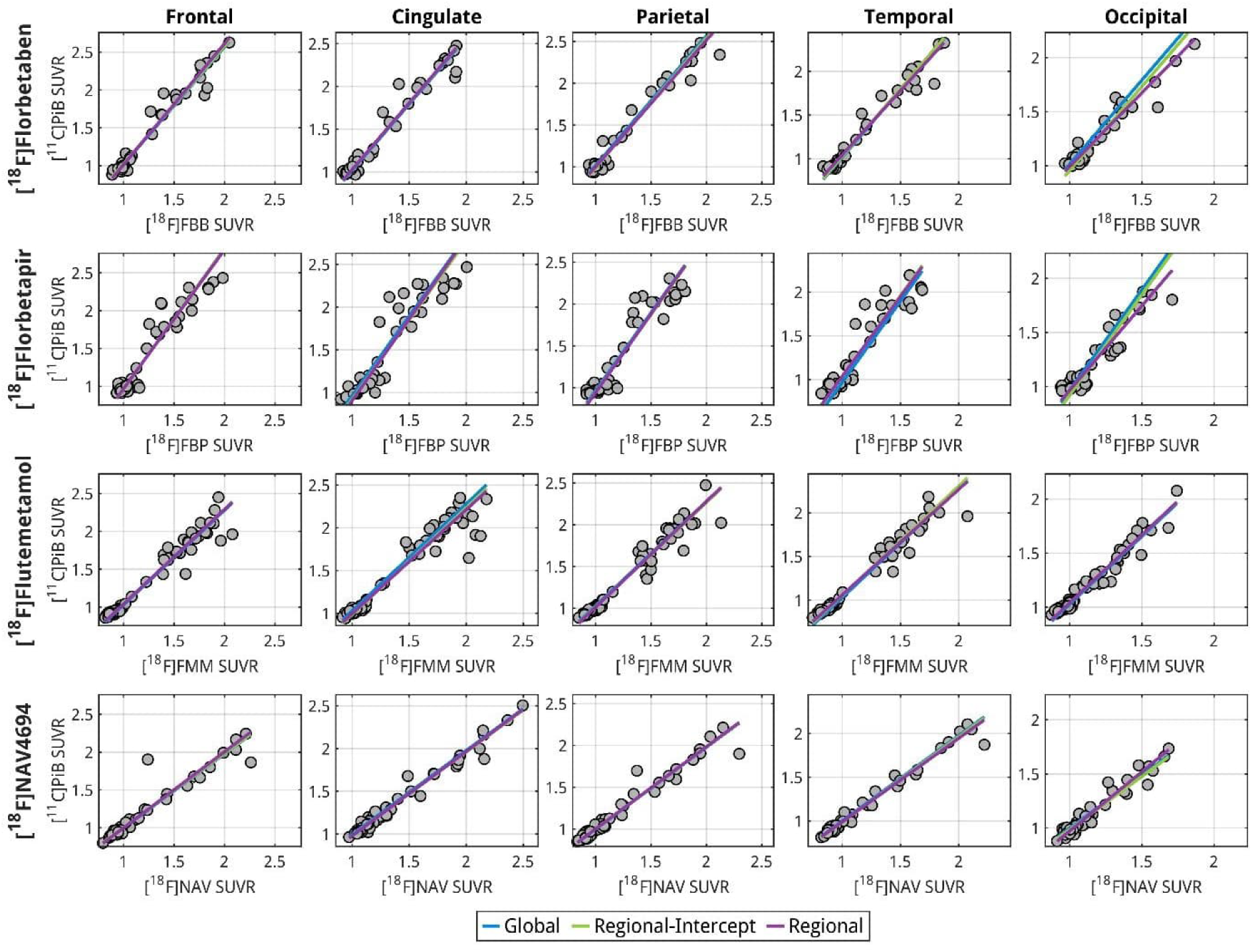
Head-to-head SUVR scatterplots and linear model fits for cortical cross-tracer harmonization. Individual regional Standardized Uptake Value Ratios (SUVR) from the head-to-head harmonization cohorts (gray circles) are plotted across cortical composite regions for [^18^F]florbetaben (FBB), [^18^F]florbetapir ([^18^F]FBP), [^18^F]flutemetamol ([^18^F]FMM), and [^18^F]NAV4694 ([^18^F]NAV) relative to reference [^11^C]PiB. Overlay lines contrast the trajectories of three evaluated calibration frameworks: the Global framework (blue), the Regional-Intercept model featuring a shared global slope and region-specific intercepts (green), and the Regional model featuring independent unconstrained slopes and intercepts per structure (purple).

**Figure 3.**
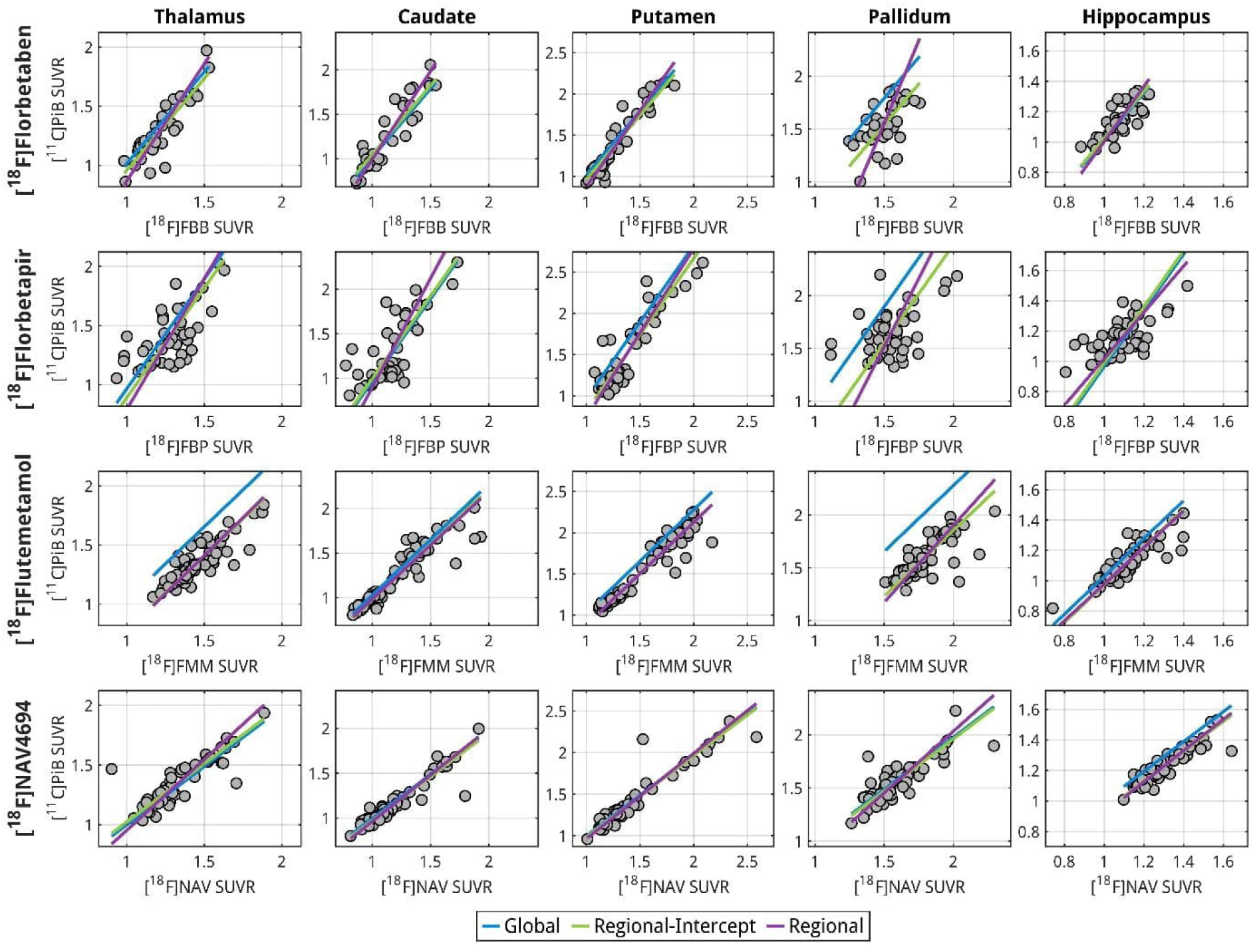
Head-to-head SUVR scatterplots and linear model fits for subcortical cross-tracer harmonization. Individual regional Standardized Uptake Value Ratios (SUVR) from the head-to-head harmonization cohorts (gray circles) are plotted across deep gray matter regions for [^18^F]florbetaben (FBB), [^18^F]florbetapir ([^18^F]FBP), [^18^F]flutemetamol ([^18^F]FMM), and [^18^F]NAV4694 ([^18^F]NAV) relative to reference [^11^C]PiB. Overlay lines contrast the trajectories of three evaluated calibration frameworks: the Global framework (blue), the Regional-Intercept model featuring a shared global slope and region-specific intercepts (green), and the Regional model featuring independent unconstrained slopes and intercepts per structure (purple).

In deep gray matter structures, including the thalamus, caudate, putamen, and pallidum, the Global regression line showed systematic offsets, failing to align the predicted [^11^C]PiB-equivalent SUVR with observed SUVR values (**Figure 3**). The Global model consistently ran above empirical data points, generating a clear bias across tracers. This overestimation peaked in the pallidum, where the Global model produced a positive bias with a MAE of 27.7 CL for [^18^F]FBB, 42.7.6 CL for [^18^F]FBP, and 45.6 CL for [^18^F]FMM, while [^18^F]NAV exhibited minimal bias (8.4 CL). The Regional-Intercept model shifted downward to align closely with observed data, eliminating this systematic offset and yielding residual conversion errors (MAE) of 15.2 CL for [^18^F]FBB, 23.7 CL for [^18^F]FBP, 8.1 CL for [^18^F]FMM, and 8.4 CL for [^18^F]NAV (**Figure 4**).

**Figure 4.**
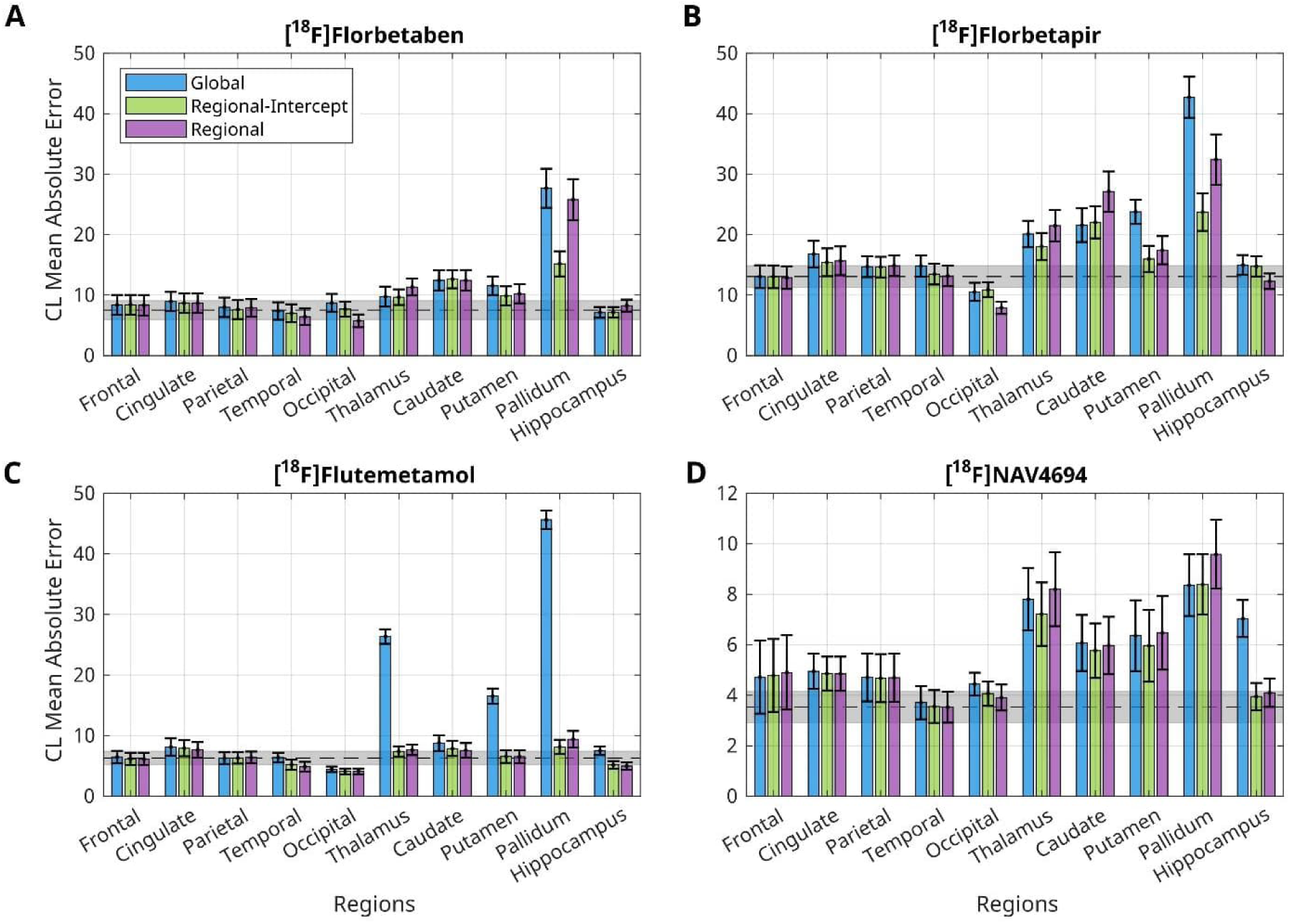
Regional Centiloid Mean Absolute Error profiles across four fluorinated tracers head-to-head dataset. The plots illustrate the regional Mean Absolute Error (*MAE_r_*) remaining after cross-tracer calibration to reference [11*C*]*PiB* and Centiloid conversion using three distinct linear mapping frameworks: Global scaling (blue), the proposed Regional-Intercept model (green), and the Regional model (purple). Panels represent evaluations for (A) [^18^F]florbetaben, (B) [^18^F]florbetapir, (C) [^18^F]flutemetamol, and (D) [^18^F]NAV4694. Colored bars indicate region-specific MAE (±standard error). The horizontal, gray-shaded band represents the native MAE (±standard error) computed across the standard global cortical composite target region to serve as a comparative benchmark. To maximize visual clarity across different tracer dynamics, Panels A through C utilize a standardized vertical axis range of 0 to 50 CL whereas Panel D retains a narrower range of 0 to 12 CL to reflect the lower baseline error variance characteristic of [^18^F]NAV4694.

These visual observations are confirmed by regional Centiloid mean absolute error profiles (**Figure 4**). The Regional-Intercept model corrected subcortical offsets on average, reducing conversion errors below 15 CL and approaching or outperforming the benchmark error of the global cortical composite. The magnitude of error reduction achieved by regional calibration was tracer-dependent. Relative to the Global model, the Regional-Intercept model reduced dataset-level error by over 50% for [^18^F]FMM (with MAE decreasing from 13.7 to 6.5 CL), while yielding moderate reductions of approximately 15% for [^18^F]FBB and [^18^F]FBP. In contrast, [^18^F]NAV4694 showed only an 8.6% improvement (from 5.8 to 5.3 CL). Among radiotracers, [^18^F]FBP exhibited the highest overall residual error across both cortical and subcortical regions **(Figure 4B**), where the fully Regional model displayed higher MAE than the Regional-Intercept model due to noise fitting and slope attenuation.

### 3.4 Voxel-Wise Topographical Centiloid Mapping

Linear conversion parameters were estimated at each voxel using the Regional-Intercept approach to generate subject-level and group-averaged voxel-wise Centiloid maps for both elderly and young control cohorts across all four fluorinated datasets and [^11^C]PiB (**Figure 5**).

**Figure 5.**
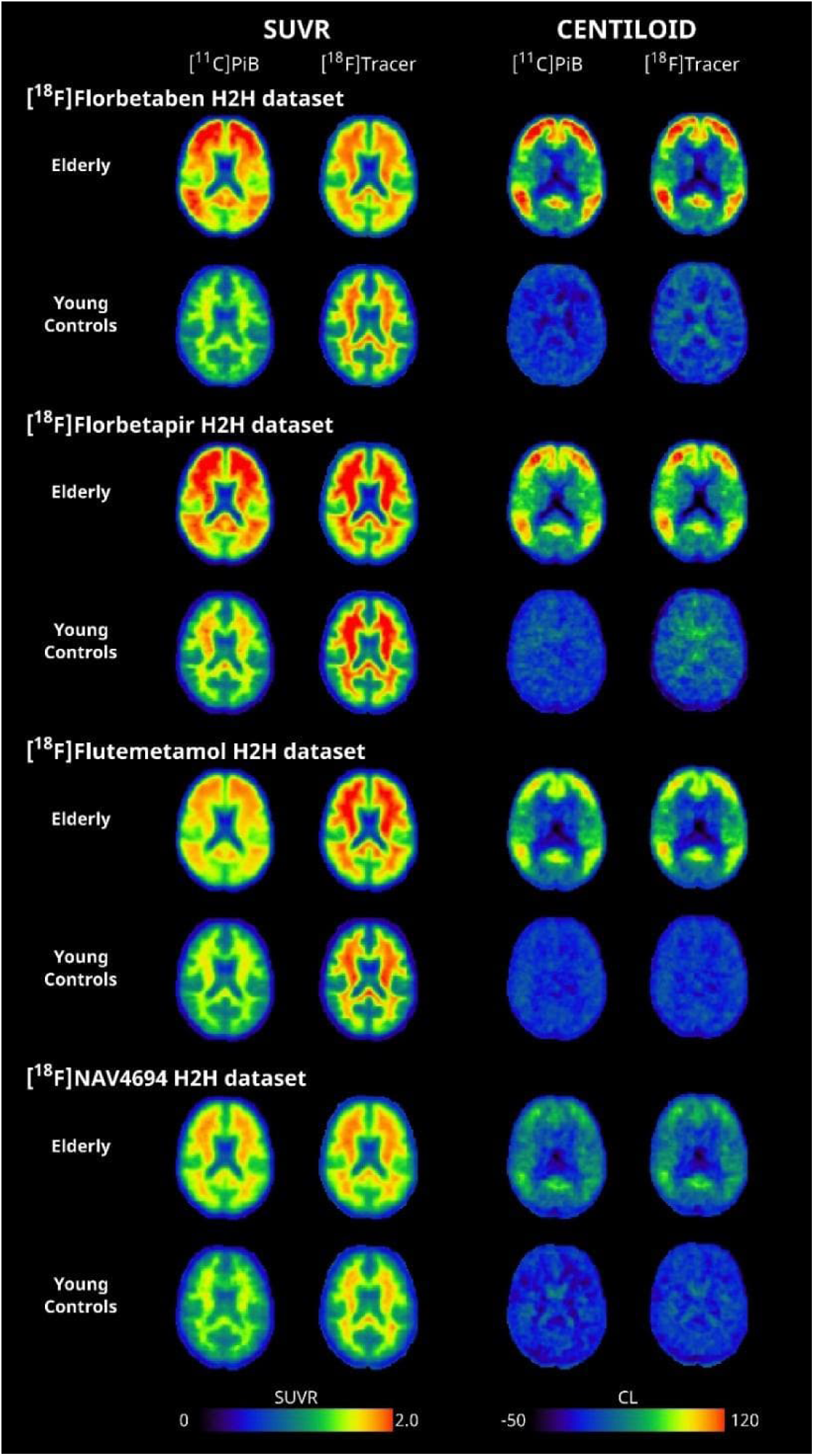
Representative voxel-wise group-averaged SUVR and Centiloid maps across fluorinated PET radiotracers. Voxel-wise group-averaged parametric maps are displayed on both the native Standardized Uptake Value Ratio scale (SUVR, left columns) and the harmonized Centiloid scale (CL, right columns) for elderly participants (top rows) and amyloid-negative young controls (YC, bottom rows) across four head-to-head fluorinated tracer datasets ([^18^F]florbetaben, [^18^F]florbetapir, [^18^F]flutemetamol, and [^18^F]NAV4694). For each tracer dataset, axial slices display the reference [^11^C]PiB acquisition side-by-side with the paired fluorinated acquisition. Centiloid parametric maps (right) were derived using the proposed Level-1 regional zero-anchor recalibration and Level-2 Regional-Intercept calibration models. Color bars represent native SUVR values (0 to 2.0) and standardized Centiloid values (−50 to 120 CL). While within-cohort concordance across tracer pairs is high, direct comparisons of absolute Centiloid burden across the four distinct elderly cohorts are precluded by differences in their clinical and diagnostic compositions, including varying proportions of mild cognitive impairment and Alzheimer disease dementia cases (Table 1).

When scaled to a native SUVR range of 0 to 2.0 (**Figure 5**, columns 1 and 2), uncalibrated images showed marked, tracer-dependent differences in spatial patterns and dynamic range driven by differential non-specific binding and dynamic ranges of fluorinated agents compared to [^11^C]PiB (Landau *et al*., 2014; Lowe *et al*., 2017; Rowe *et al*., 2017; Chen *et al*., 2023). While [^11^C]PiB and [^18^F]NAV4694 displayed minimal non-displaceable white matter retention and sharp target-to-background contrast, [^18^F]florbetaben and [^18^F]florbetapir exhibited high non-specific white matter binding and narrowed dynamic ranges. These regional variations in non-displaceable uptake prevented direct cross-tracer comparisons. Centiloid conversion improved spatial concordance and cross-tracer consistency within each cohort (**Figure 5**, columns 3 and 4). Neocortical amyloid retention patterns converged between paired [^11^C]PiB and fluorinated scans, with key regions of early accumulation, such as the precuneus, posterior cingulate, and frontal cortices, exhibiting clear demarcation from surrounding background tissues. While non-specific white matter signal was substantially reduced, residual differences in subcortical white matter remained evident between tracers. Consistent with this visual convergence, mutual information between [¹¹C]PiB and fluorinated maps increased across all elderly cohorts from SUVR to Centiloid space ([^18^F]florbetaben: 1.15 to 1.58; [^18^F]florbetapir: 1.12 to 1.54; [^18^F]flutemetamol: 1.47 to 1.77; [^18^F]NAV4694: 1.54 to 1.66). Importantly, although we observed high within-cohort concordance across tracer pairs, direct comparisons of absolute Centiloid burden across the four distinct elderly cohorts are precluded by differences in their clinical and diagnostic compositions, such as varying proportions of MCI and AD dementia cases (**Table 1**).

## 4. Discussion

Standard global Centiloid metrics remain the benchmark for defining amyloid positivity and clinical trial inclusion criteria. However, regional measures are increasingly needed to detect early preclinical stages and to track amyloid-lowering therapies, because a global average can hide focal plaque accumulation and uneven clearance. Here, we developed a standardized regional and voxel-wise Centiloid framework across four fluorinated tracers ([^18^F]florbetaben, [^18^F]florbetapir, [^18^F]flutemetamol, and [^18^F]NAV4694) to complement standard global metrics. In short, the global equations work well in the neocortex but cause large errors in deep gray matter, and adding region-specific intercepts corrects these errors.

### 4.1 Regional Centiloid Framework

#### 4.1.1 Level-1 Regional Scale Anchoring

At Level-1, non-specific tracer uptake in young controls differed across brain regions (**Figure 1A**), reflecting non-displaceable signal rather than amyloid pathology. Anchoring each region to its young control average using a region-specific intercept centered all ten structures at 0 CL (Figure 1B), preserving a consistent biological definition of 0 CL as the absence of amyloid in every region.

#### 4.1.2 Level-2 Model Selection and Regional-Intercept Calibration

At Level-2, the Regional-Intercept model was selected as optimal for all four radiotracers (**Table 2**, **Figure 4**). Applying standard global equations directly to individual structures introduced substantial systematic error, particularly in deep gray matter. On the other hand, fitting a separate slope and intercept for each region (the fully Regional model) was counterproductive in regions with low correlation, because inverting an unstable regional slope magnifies measurement noise. Keeping a shared global slope while adjusting only the regional intercepts corrects the regional offsets without amplifying noise, maintaining subcortical MAE on average below 15 CL.

The performance of the global equations depends on the brain area: they work well in the neocortex but are biased in deep subcortical structures. In the main neocortical lobes, the Global and Regional-Intercept fits nearly overlap with the fully Regional fit (**Figure 2**). These regions have a dynamic range similar to the composite, so the global calibration is highly accurate there. The occipital cortex, which is not part of the standard cortical target region, is an exception: here, the Regional-Intercept model eliminates systematic inter-model offsets (up to 6.4 CL, **Figure 2**) and removes directional bias, while yielding modest changes in overall conversion error relative to the global model (**Figure 4**). In deep gray matter, by contrast, applying the global equations directly introduces substantial systematic error, reaching 27.7–45.6 CL in the pallidum (Figures 3 and 4). This likely explains the implausibly high subcortical values reported previously (Noh *et al*., 2026). These biases reflect differences in subcortical non-displaceable binding between [^11^C]PiB and fluorinated tracers (Landau *et al*., 2014; Rowe *et al*., 2017), creating regional variations that a single global equation cannot capture. Among the tracers, [^18^F]florbetapir had the highest MAE, possibly because of its shorter 10-minute acquisition window and different processing (**Figure 4B**). Even so, the Regional-Intercept model remained the best model in all cohorts.

#### 4.1.3 Same Centiloid Value, Same Amyloid Concentration

Combining the Level-1 zero anchoring with the Level-2 Regional-Intercept calibration gives a scale on which the same Centiloid value reflects the equivalent amyloid concentration, in any region and with any tracer. For example, the same value in the pallidum and in the frontal cortex corresponds to the same amount of amyloid. Keeping a shared global slope respects the fact that regions differ in how much amyloid they can accumulate (Whittington, Sharp and Gunn, 2018), instead of artificially stretching each region between 0 and 100 CL. This has a practical advantage for longitudinal studies. Because the Global and Regional-Intercept models share the same slope, the regional intercepts cancel out when a baseline value is subtracted from a follow-up value in the same person. Therefore, both models give the same change over time, and the standard global equations remain suitable for measuring regional longitudinal changes, such as plaque clearance during treatment.

#### 4.1.4 Voxel-Wise Centiloid Maps

At the voxel level, the Regional-Intercept conversion substantially improved spatial agreement across all fluorinated tracers (**Figure 5**). In elderly cohorts, cortical retention matched known neuropathological patterns in early-accumulating regions (Thal *et al*., 2002; Grothe *et al*., 2017), and mutual information with [^11^C]PiB increased. Some cross-tracer differences remained in deep white matter. This is expected, because the calibration targeted gray matter, and our pipeline does not apply partial volume correction (to avoid amplifying noise). As a result, signal spillover between white matter and the adjacent cortex is not corrected, leaving residual cross-tracer variation in deep white matter.

### 4.2 Potential Clinical Uses

Standard global Centiloid metrics are essential for stratifying trial participants and for defining treatment-related amyloid clearance thresholds in clinical trials (La Joie *et al*., 2025).

Under the standard global equations, uncalibrated non-specific signal can inflate subcortical values beyond 190 CL before treatment (Noh *et al*., 2026). This shows the need for regional scale anchoring. Because the regional intercepts cancel out when computing changes over time (see above), regional calibration is mainly useful for measuring burden at a single time point and for setting regional positivity thresholds. Beyond these direct corrections, our framework may support other applications. First, it may help define regional negativity, for example as a criterion to stop treatment. Because every region shares the same zero and the same slope, an investigator could in principle apply the same Centiloid threshold to the whole brain. However, caution is warranted: regional differences in measurement noise, tissue volume, and uncorrected partial volume effects maintain structure-specific variance. Establishing definitive regional negativity thresholds might therefore require calibration against histology.

Second, regional Centiloid metrics may clarify how amyloid interacts with downstream pathology in specific regions. Neocortical tau-PET signal rises sharply once global amyloid reaches 40–70 CL (Collij *et al*., 2024). In trials, clinical benefit differed depending on initial tau burden (Sims *et al*., 2023; Chiotis *et al*., 2026), and interactions between tau burden and amyloid removal have been reported (Tonietto et al. AAIC2026 presentation). Regional amyloid metrics may therefore help identify the best time window for treatment, before tau spreads widely.

Finally, regional measures may also help with diagnosis. Visual reads and quantitative values disagree in the intermediate 10-40 CL range in approximately 14% of real-world cases (Zeltzer *et al*., 2025) and 11% of clinical memory clinic cohorts (Collij *et al*., 2024). In early disease, focal plaques in structures like the precuneus or posterior cingulate can be clearly visible but not enough to raise the global composite above the positivity threshold. Regional values and voxel-wise maps (**Figure 5**) could help bridge visual reads and quantitative Centiloid values. Whether they actually reduce variability between readers or improve the classification of borderline cases is still untested and requires dedicated clinical reader studies.

### 4.3 Limitations

Most technical, demographic, and cohort-size constraints come from the original Centiloid standard and are not introduced by our method. Still, some limitations should be considered. First, the standard calibration datasets are small (25–50 elderly participants; 10–24 YC per tracer). These data are the established benchmark for cross-tracer calibration, but the regional conversion parameters should be confirmed in larger head-to-head cohorts and clinical trials (Zhang *et al*., 2025). Longitudinal studies should also confirm that the parameters remain stable during active plaque clearance, when structural changes or amyloid-related imaging abnormalities alter local brain anatomy (La Joie *et al*., 2025). Second, PET hardware and image processing introduce variability. The calibration scans were acquired on earlier PET systems. Modern digital scanners offer higher resolution, better signal-to-noise ratio, and sharper gray-to-white matter boundaries, which may affect region segmentation and partial volume effects. Partial volume correction (PVC) was not applied, because it requires scanner-specific tuning, which prevents a unified workflow, and it amplifies noise. Because PVC changes the tracer dynamic range and the non-displaceable background, our non-PVC calibration parameters cannot be applied to PVC-corrected data, which would require independent calibrations. In addition, regional values also depend on structural MRI processing, which is sensitive to registration errors in severe atrophy (Markiewicz *et al*., 2021; Bourgeat *et al*., 2026). Finally, the standard datasets lack racial and ethnic diversity (Collij *et al*., 2024), highlighting the need for further validation in representative global cohorts (Wilkins *et al*., 2022).

### 4.4 Conclusions

This study establishes a Regional-Intercept Centiloid framework extending standardized amyloid-PET quantification to regional and voxel-wise scales. For clinical and translational researchers, our findings provide four direct takeaways:

1. Global equations work in the neocortex: In most neocortical gray matter, the standard global equations give comparable Centiloid values, as confirmed across the major lobes.

1. Deep gray matter requires regional intercept calibration: The standard global equations systematically overestimate subcortical values, with conversion errors reaching a MAE of 45.6 CL in the pallidum. Adding region-specific intercepts removes this systematic offset, although some structure-specific residual variability remains around the harmonized 0 CL.
2. Tracking change over time: The standard global equations reliably track longitudinal change and therapy response in all structures, because the regional offsets cancel out when values are subtracted. However, defining meaningful regional negativity thresholds remains an open question that might require histological calibration.
3. Same Centiloid value, same amyloid concentration: With a shared zero point and a single slope, the same Centiloid value corresponds approximately to the same amyloid concentration across regions and tracers. The equivalence is not exact, because partial volume effects are not corrected, but this approach supports multi-center pooling of regional amyloid measurements across radiotracers.

As disease-modifying therapies enter clinical practice, standardized regional metrics provide the anatomical detail needed to evaluate patchy clearance, detect early focal accumulation, and study how amyloid interacts with tau pathology in specific regions.

## Author Contributions

**Erica Silvestri:** Conceptualization; formal analysis; methodology; visualization; writing–original draft. **Pavanjit Chaggar:** Formal analysis; Methodology, writing–review and editing. **Susanna Gobbi:** Formal analysis; Methodology, writing–review and editing. **Roxana Aldea:** Methodology; writing–review and editing. **Gregory Klein:** Conceptualization; methodology; writing–review and editing. **Matteo Tonietto:** Conceptualization; methodology; supervision; writing–review and editing.

## Ethics Approval and Consent to Participate

Primary PET and MRI data used in this study were obtained from the open-access Global Alzheimer’s Association Interactive Network (GAAIN) Centiloid Project repository (www.gaain.org/centiloid-project). All original reference cohorts included in the Level-1 and Level-2 Centiloid calibration datasets were acquired in accordance with the ethical principles of the Declaration of Helsinki. Original study protocols were approved by the local Institutional Review Boards and Human Research Ethics Committees of the primary contributing centers (including the University of Pittsburgh, Austin Health Melbourne, Mayo Clinic, and Avid Radiopharmaceuticals), and all participants or their legal representatives provided written informed consent prior to data collection.

## Data and Code Availability

Primary PET and MRI datasets analyzed in this study are publicly available through the Global Alzheimer’s Association Interactive Network (GAAIN) Centiloid Project repository (https://www.gaain.org/centiloid-project). All derived calibration parameters, region-specific conversion equations, and atlas lookup values are included in the Supplementary Materials (Supplementary Tables S3–S8). The complete Python processing code, Regional-Intercept implementation scripts, and precomputed MNI152 standard-space parameter maps are publicly hosted on https://github.com/Erica-Silvestri/AmyloidPET-Regional-Centiloid-Conversion.git

