## Supplementary Materials for "Harmonizing amyloid PET tracers at the local level: evaluation of regional Centiloid equations"

### Supplemental Online Content

#### Supplementary Methods

##### 1. Image Processing

Image processing and quantification followed the Alzheimer's Disease Neuroimaging Initiative native-space PET pipeline (Royse *et al.*, 2021). Structural T1w MRI scans were processed using FreeSurfer *recon-all* (v6.0, (Fischl, 2012)) for automated parcellation based on the Desikan-Killiany atlas for cortical regions and the probabilistic Gaussian Classifier Atlas for subcortical structures. Structural images were spatially normalized to FSL MNI152 standard space using non-linear diffeomorphic registration via Advanced Normalization Tools (ANTs, (Avants *et al.*, 2011)).
Time-averaged PET volumes were rigidly registered to the corresponding T1w structural image using FreeSurfer *mri_coreg*. Regional and global target Standardized Uptake Value Ratios (SUVRs) were calculated using the whole cerebellum as the reference region. For voxel-wise analyses, time-averaged PET images were mapped into MNI152 2 mm standard space by concatenating the rigid PET-to-T1w transform with the T1w-to-MNI non-linear warp in a single interpolation step. No additional spatial smoothing or image-based filtering was applied.

##### 2. PET Preprocessing and Acquisition Windows

For the [$^{18}$F]florbetapir head-to-head dataset, PET images were preprocessed using SPM12 (Statistical Parametric Mapping) for motion correction and temporal frame averaging. All dynamic PET acquisitions were realigned to correct for subject motion using the default SPM12 rigid body realignment algorithm. Time-averaged volumes corresponding to the tracer-specific acquisition windows (50–60 min for [$^{18}$F]florbetapir; 50–70 min for [$^{11}$C]PiB) were generated by averaging individual motion-corrected frames. For all other Level-1 anchoring and Level-2 head-to-head datasets ([$^{18}$F]florbetaben, [$^{18}$F]flutemetamol, [$^{18}$F]NAV4694), PET preprocessing was performed by the GAAIN consortium following standardized protocols (Klunk et al., 2015). Pre-processed time-averaged volumes corresponding to the tracer-specific acquisition windows were obtained directly from the GAAIN data repository.
No additional spatial smoothing or image-based filtering was applied to preserve regional quantification accuracy.

##### 3. Regions of Interest Definitions

Table S1 provides a comprehensive list of the ten primary regions of interest (ROIs) evaluated in this study, including cortical lobes (frontal, cingulate, parietal, temporal, occipital) and subcortical deep gray matter structures (thalamus, caudate, putamen, pallidum, hippocampus), alongside the standard ADNI global cortical composite target region. Cortical regions of interest were defined using the Desikan-Killiany atlas, whereas subcortical structures were segmented using the probabilistic Gaussian Classifier Atlas, both derived from the automated whole-brain segmentation stream implemented in FreeSurfer (v6.0; (Fischl, 2012)). For fine-grained analyses, individual bilateral Desikan-Killiany cortical subregions were also extracted as detailed below under Single Desikan-Killiany Atlas Region Implementation.

##### 4. Data Imputation for Level-1 Reference Cohort

During structural MRI preprocessing quality control, two Alzheimer's disease participants in the Level-1 [$^{11}$C]PiB anchoring dataset were excluded from regional parcellation analysis due to severe volumetric segmentation failures in FreeSurfer (global brain atrophy with poor gray-white matter boundary definition). To maintain the exact sample size required for standard GAAIN global equation recalibration (N=45 AD participants), global whole-cerebellum reference SUVR values for these two excluded participants were mathematically imputed using whole-group mean substitution based on diagnostic classification (AD participants).

Importantly, the proposed regional equations utilize Level-1 data strictly to establish the regional zero-anchor point using the young control cohort (N=34 unaffected). Therefore, this AD-level imputation carries no mathematical weight in deriving regional conversion formulas.

### 5. Fine-Grained Regional Conversion: From Composite Regions to Individual Atlas Regions

Following the identification of the Regional-Intercept model as the optimal calibration framework (main Results, Section 3.3), we extended the regional conversion approach from the ten primary composite ROIs to individual bilateral Desikan-Killiany (DK) atlas regions for enhanced anatomical specificity.

#### 5.1. Single Desikan-Killiany Atlas Region Implementation

For each individual bilateral cortical region, region-specific intercepts were determined while preserving the global slope derived from Level-2 calibration. To enhance the signal-to-noise ratio of SUVR measurements and mitigate statistical variance in smaller structures, left and right hemispheric values were averaged prior to equation estimation:

$$SUVR_{bilateral} = \frac{SUVR_{left} + SUVR_{right}}{2}$$

Regional Centiloid values were then computed as:

$$CL_{region} = \left(\frac{m_{PiB}}{m_F}\right) \cdot SUVR_{F,\, bilateral} + B_{CL,\, region}$$

where:

- $m_{PiB}$ is the global [$^{11}$C]PiB scaling slope (107.67)
- $m_F$ is the global conversion slope derived from Level-2 head-to-head calibration
- $SUVR_{F,\, bilateral}$ is the averaged fluorinated tracer uptake across left and right hemispheres
- $B_{CL,\, region}$ is the region-specific Centiloid intercept, pre-integrating the Level-2 head-to-head offset with the Level-1 young control zero-anchor point

Sensitivity analyses comparing left-hemisphere, right-hemisphere, and bilateral-averaged approaches showed minimal differences in final Centiloid values (mean absolute difference $< 2\ CL$), confirming that bilateral averaging preserves regional accuracy while improving signal stability.

#### 5.2. Voxel-Wise Conversion Framework

To generate continuous voxel-wise parametric maps, the Regional-Intercept calibration framework was implemented at the voxel level across standard MNI152 space (2 mm³ resolution). This pipeline integrates Level-1 scale anchoring with Level-2 cross-tracer calibration into a precomputed voxel-wise conversion map:

<u>Level-1 Scale Anchoring:</u>

At each voxel $v$, the biological zero-anchor point was defined as the mean [$^{11}$C]PiB uptake across the young control cohort scans in standard space:

$$Anchor_{0,v} = \underline{SUVR}_{PiB,\, YC,v}$$

Using the standard global [$^{11}$C]PiB scaling slope ($m_{PiB}$ = 107.67), the voxel-specific [$^{11}$C]PiB intercept ($b_{PiB,v}$) was established as:

$$b_{PiB,v} = -m_{PiB} \cdot Anchor_{0,v}$$

This anchors [11C]PiB Centiloid values to center at $0\ CL$ across all brain voxels in young controls:

$$CL_{PiB,v} = m_{PiB} \cdot SUVR_{PiB,v} + b_{PiB,v} = m_{PiB} \cdot \left(SUVR_{PiB,v} - Anchor_{0,v}\right)$$

Level-2 Cross-Tracer Calibration:

For each fluorinated tracer, the global conversion gradient ($m_F$, regressing fluorinated SUVR onto [11C]PiB SUVR) was fixed from the global Level-2 calibration to maintain scale stability. At each voxel $v$, the local tracer intercept ($b_{F,v}$) was calculated across the paired head-to-head dataset by taking the mean difference between observed fluorinated uptake and gradient-scaled $[11C]PiB$ uptake:

$$b_{F,v} = \frac{1}{N}\sum_{i=1}^{N} \left(SUVR_{F,i,v} - m_F \cdot SUVR_{PiB,i,v}\right)$$

Inverting this relationship yields the predicted [11C]PiB -equivalent SUVR at voxel $v$:

$$\widehat{SUVR}_{PiB,v} = \frac{SUVR_{F,v} - b_{F,v}}{m_F}$$

Direct Voxel-Wise Centiloid Transformation:

Substituting predicted $[11C]PiB$ SUVR into the Level-1 equation defines the final Centiloid value at voxel $v$:

$$CL_{F,v} = m_{PiB} \cdot \left(\frac{SUVR_{F,v} - b_{F,v}}{m_F} - Anchor_{0,v}\right)$$

In practice, this formulation simplifies into a single precomputed parametric map:

$$CL_{F,v} = \left(\frac{m_{PiB}}{m_F}\right) \cdot SUVR_{F,v} + B_{CL,v}$$

where the composite voxel-wise Centiloid intercept map ($B_{CL,v}$) pre-integrates both calibration levels:

$$B_{CL,v} = -\left(\frac{m_{PiB}}{m_F}\right) \cdot b_{F,v} + b_{PiB,v}$$

This formulation eliminates artificial subcortical offsets and local non-displaceable background variations at every voxel while preserving the robust global dynamic range across the entire brain volume.

Here, $b_{F,v}$ resolves local tracer-specific non-displaceable differences derived from the head-to-head paired scans, while $b_{PiB,v}$ anchors the zero CL point using the young control cohort.

Calculations were restricted to the intracranial brain volume using the dilated MNI152 brain mask to prevent extra-cerebral background artifacts.

#### 5.3. Voxel-Wise Mutual Information Calculation

To quantify spatial concordance and cross-tracer alignment in voxel space, mutual information was calculated using Advanced Normalization Tools (ANTs) via the Mattes mutual information metric. Calculations were performed on the group-averaged maps within each elderly cohort, comparing paired $[11C]PiB$ against each fluorinated tracer before and after Centiloid conversion. To prevent potential biases from extra-cerebral signals, air voxels, and zero-padding artifacts, computation was strictly confined to the standard brain-extracted mask of the MNI152 2 mm template provided by FSL. Mattes mutual information was calculated using 32 joint histogram bins with probability density estimation normalized over the masked voxels. Because binning is normalized over the distribution within the mask, the resulting values reflect shared spatial entropy and shared anatomical retention patterns rather than artificial scaling artifacts arising from differences in numerical dynamic range between native SUVR and Centiloid units.

### Table S1. GAAIN Centiloid Calibration Specifications & Cohort DemographicsTable S2. List of Regions of Interests

| Specification Category | Level-1 [$^{11}$C]PiB (Anchoring Scale) | Level-2 [$^{18}$F]Florbetaben (FBB / Neuraceq) | Level-2 [$^{18}$F]Florbetapir (FBP / Amyvid) | Level-2 [$^{18}$F]Flutemetamol (FMM / Vizamyl) | Level-2 [$^{18}$F]NAV4694 (NAV / Flutafuranol) |
|---|---|---|---|---|---|
| **Primary GAAIN Center & Key Reference** | University of Pittsburgh<br>(Klunk et al., 2015) | Austin Health, Melbourne<br>(Rowe *et al.*, 2017) | Avid / Multi-Center<br>(Navitsky *et al.*, 2018) | GE Healthcare / Mayo Clinic<br>(Battle *et al.*, 2018) | Austin Health, Melbourne<br>(Rowe *et al.*, 2016) |
| **PET Scanner Models** | Siemens ECAT EXACT HR+;<br>GE Advance | Philips Allegro (N=6);<br>Philips TF64 PET/CT (N=29) | Siemens ECAT EXACT HR+;<br>Philips Gemini TF 64;<br>GE Advance | GE Discovery 710 / 690;<br>Philips Gemini TF | Philips Allegro PET system |
| **Detector Crystal Types** | BGO / NaI | BGO (Allegro);<br>LYSO (TF64) | BGO (HR+, Advance);<br>LYSO (Gemini TF) | LYSO / BGO | BGO |
| **MRI Hardware Specifications** | 1.5T / 3T Siemens / GE | 3T Siemens Magnetom Trio / Prisma | 3T Siemens / GE / Philips | 1.5T / 3T MRI systems | 3T Siemens Magnetom Trio |
| **Raw PET Matrix & Voxel Size** | 128x128 / 256x256<br>2.0x2.0x2.0 mm³ isotropic | 128x128 / 256x256<br>2.0x2.0x2.0 mm³ isotropic | 128x128 / 256x256<br>2.0 to 2.34 mm isotropic | 128x128 / 256x256<br>2.0 to 2.34 mm isotropic | 128x128 / 256x256<br>2.0x2.0x2.0 mm³ isotropic |
| **Reconstruction Algorithms** | 2D/3D OSEM, FORE-iterative | 3D LOR-RAMLA /<br>FORE-iterative | 2D-OSEM (HR+), LOR-RAMLA,<br>FORE-iterative (Advance) | 3D-OSEM<br>(4 iterations, 18 subsets) | 3D Row-Action Maximum Likelihood (RAMLA) |
| **Total Cohort Sample Size (N)** | N = 79<br>(34 YC, 45 AD) | N = 35<br>(10 YC, 25 Elderly) | N = 46<br>(13 YC, 33 Elderly) | N = 74<br>(24 YC, 50 Elderly) | N = 55<br>(10 YC, 45 Elderly) |
| **Young Controls (YC / YHC) [N, Age, Sex, MMSE]** | N = 34<br>• Age: 24 ± 3 yr<br>• Sex: 18M / 16F<br>• MMSE: 29.8 ± 0.5 | N = 10<br>• Age: 33 ± 8 yr (<45y)<br>• Sex: 5M / 5F<br>• MMSE: >28 | N = 13<br>• Age: 30 ± 6 yr (<45y)<br>• Sex: 7M / 6F<br>• MMSE: ≥29 | N = 24<br>• Age: 37 ± 5 yr (30−45y)<br>• Sex: 10M / 14F<br>• MMSE: ≥29 | N = 10<br>• Age: 33 ± 7 yr (<45y)<br>• Sex: 4M / 6F<br>• MMSE: ≥28 |
| **Elderly Healthy Controls (HC) [N, Age, Sex, MMSE]** | N/A | N = 6<br>• Age: 71.3 ± 8.0 yr<br>• Sex: 3M / 3F<br>• MMSE: 29 ± 1 | N = 9<br>• Age: 74 ± 7 yr<br>• Sex: 5M / 4F<br>• MMSE: 29 ± 1 | N = 10<br>• Age: 57 ± 11 yr<br>• Sex: 4M / 6F<br>• MMSE: 29 ± 1 | N = 25<br>• Age: 74 ± 8 yr<br>• Sex: 11M / 14F<br>• MMSE: 29 ± 1 |
| **Mild Cognitive Impairment (MCI) [N, Age, Sex, MMSE]** | N/A | N = 9<br>• Age: 72 ± 5 yr<br>• Sex: 5M / 4F<br>• MMSE: 28 ± 2 | N = 7<br>• Age: 73 ± 6 yr<br>• Sex: 4M / 3F<br>• MMSE: 27 ± 2 | N = 20<br>• Age: 73 ± 7 yr<br>• Sex: 8M / 12F<br>• MMSE: 27 ± 2 | N = 10<br>• Age: 75 ± 9 yr<br>• Sex: 6M / 4F<br>• MMSE: 27 ± 3 |
| **Alzheimer's Disease (AD) [N, Age, Sex, MMSE]** | N = 45<br>• Age: 71.5 ± 8.9 yr<br>• Sex: 22M / 23F<br>• MMSE: 21.2 ± 4.1 | N = 8<br>• Age: 69 ± 6 yr<br>• Sex: 4M / 4F<br>• MMSE: 23 ± 3 | N = 17<br>• Age: 71 ± 8 yr<br>• Sex: 9M / 8F<br>• MMSE: 21 ± 4 | N = 20<br>• Age: 69 ± 10 yr<br>• Sex: 8M / 12F<br>• MMSE: 21 ± 4 | N = 7<br>• Age: 73 ± 11 yr<br>• Sex: 3M / 4F<br>• MMSE: 24 ± 2 |
| **Frontotemporal Dementia (FTD) [N, Age, Sex, MMSE]** | N/A | N = 2<br>• Age: 74 ± 8 yr<br>• Sex: 1M / 1F<br>• MMSE: 23 ± 1 | N = 0 | N = 0 | N = 3<br>• Age: 68 ± 5 yr<br>• Sex: 2M / 1F<br>• MMSE: 27 ± 1 |
| **[11C]PiB Injected Dose & Window** | 555 ± 55 MBq (15 ± 1.5 mCi)<br>50-70 min (4x5 min frames) | 555 ± 55 MBq (15 ± 1.5 mCi)<br>50-70 min (4x5 min frames) | 370-555 MBq (10-15 mCi)<br>50-70 min (4x5 min frames) | 370-555 MBq (10-15 mCi)<br>50-70 min (4x5 min frames) | 370 ± 37 MBq (10 ± 1.0 mCi)<br>50-70 min (4x5 min frames) |
| **Fluorinated Tracer Dose & Window** | N/A | 300 ± 30 MBq (8.1 ± 0.8 mCi)<br>90-110 min (4x5 min frames) | 370 ± 37 MBq (10 ± 1.0 mCi)<br>50-60 min (2x5 min frames) | 185 ± 18.5 MBq (5.0 ± 0.5 mCi)<br>90-110 min (4x5 min frames) | 250 ± 25 MBq (6.8 ± 0.7 mCi)<br>50-70 min (4x5 min frames) |

| Region | Atlas Labels | Notes |
|---|---|---|
| Composite Target Region | Cingulate, Frontal, Parietal, Temporal | Weighted average of the composing regions (Royse *et al.*, 2021) https://adni.bitbucket.io/reference/docs/UCBERKELEYAV45/UCBERKELEY_AV45_Methods_11.15.2021.pdf |
| Cingulate | ctx-lh-caudalanteriorcingulate, ctx-lh-isthmuscingulate, ctx-lh-posteriorcingulate, ctx-lh-rostralanteriorcingulate, ctx-rh-caudalanteriorcingulate, ctx-rh-isthmuscingulate, ctx-rh-posteriorcingulate, ctx-rh-rostralanteriorcingulate | Defined as in (Klein and Tourville, 2012) |
| Frontal | ctx-lh-caudalmiddlefrontal, ctx-lh-lateralorbitofrontal, ctx-lh-medialorbitofrontal, ctx-lh-parsopercularis, ctx-lh-parsorbitalis, ctx-lh-parstriangularis, ctx-lh-rostralmiddlefrontal, ctx-lh-superiorfrontal, ctx-lh-frontalpole, ctx-rh-caudalmiddlefrontal, ctx-rh-lateralorbitofrontal, ctx-rh-medialorbitofrontal, ctx-rh-parsopercularis, ctx-rh-parsorbitalis, ctx-rh-parstriangularis, ctx-rh-rostralmiddlefrontal, ctx-rh-superiorfrontal, ctx-rh-frontalpole | Defined as *Frontal regions* in (Landau *et al.*, 2021) |
| Occipital | ctx-lh-lateraloccipital, ctx-lh-lingual, ctx-lh-cuneus, ctx-lh-pericalcarine, ctx-rh-lateraloccipital, ctx-rh-lingual, ctx-rh-cuneus, ctx-rh-pericalcarine | Defined as in (Landau *et al.*, 2021) |
| Parietal | ctx-lh-inferiorparietal, ctx-lh-precuneus, ctx-lh-superiorparietal, ctx-lh-supramarginal, ctx-rh-inferiorparietal, ctx-rh-precuneus, ctx-rh-superiorparietal, ctx-rh-supramarginal | Defined as *Lateral parietal regions* in (Landau *et al.*, 2021) |
| Temporal | ctx-lh-middletemporal, ctx-lh-superiortemporal, ctx-rh-middletemporal, ctx-rh-superiortemporal | Defined as *Lateral temporal regions* in (Landau *et al.*, 2021) |
| Caudate | Left-Caudate, Right-Caudate | |
| Pallidum | Left-Pallidum, Right-Pallidum | |
| Putamen | Left-Putamen, Right-Putamen | |
| Thalamus | Left-Thalamus-Proper, Right-Thalamus-Proper | |
| White Matter | Left-Cerebral-White-Matter, Right-Cerebral-White-Matter | The mask was eroded using a spherical kernel with a radius of 3 mm. |
| Cerebellum | Left-Cerebellum-Cortex, Right-Cerebellum-Cortex, Left-Cerebellum-White-Matter, Right-Cerebellum-White-Matter | |

### Supplementary Results

### Figure S1: Global Formula Level-1 Validation

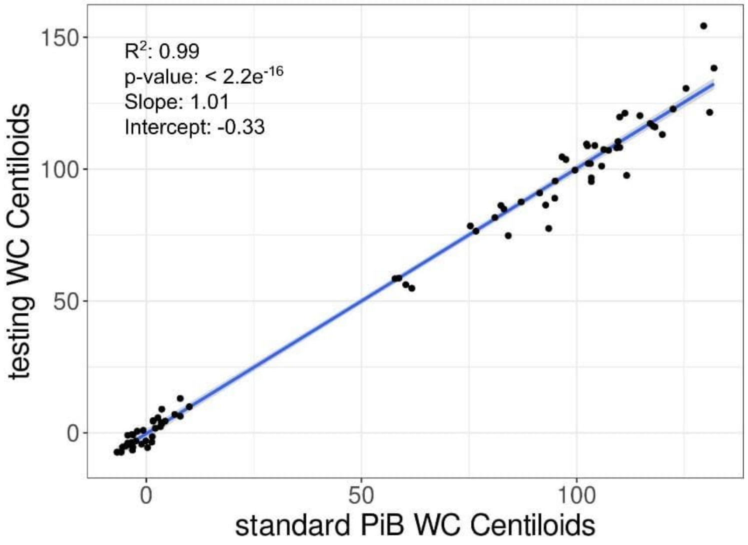


**Figure S1. Level-1 validation of the global Centiloid equation.** Scatterplot shows the correlation between Centiloid (CL) values from the standard [11C]PiB dataset provided by GAAIN (x-axis) and the CL values obtained from the same dataset using our internal pipeline (y-axis). The regression analysis indicates strong concordance with a coefficient of determination ($R^2$) of 0.99, a slope of 1.01, and an intercept of -0.33, confirming that all established Centiloid Level-1 validation criteria were fully met (slope $1.00 \pm 0.02$, intercept $0.00 \pm 2.00$, and $R^2 > 0.98$). The blue line represents the linear fit, with the gray shaded area indicating the 95% confidence interval. (p-value < 2.2 10-16). WC = Whole Cerebellum reference region.

**Figure S2. Global Formula Level-2 Validations**

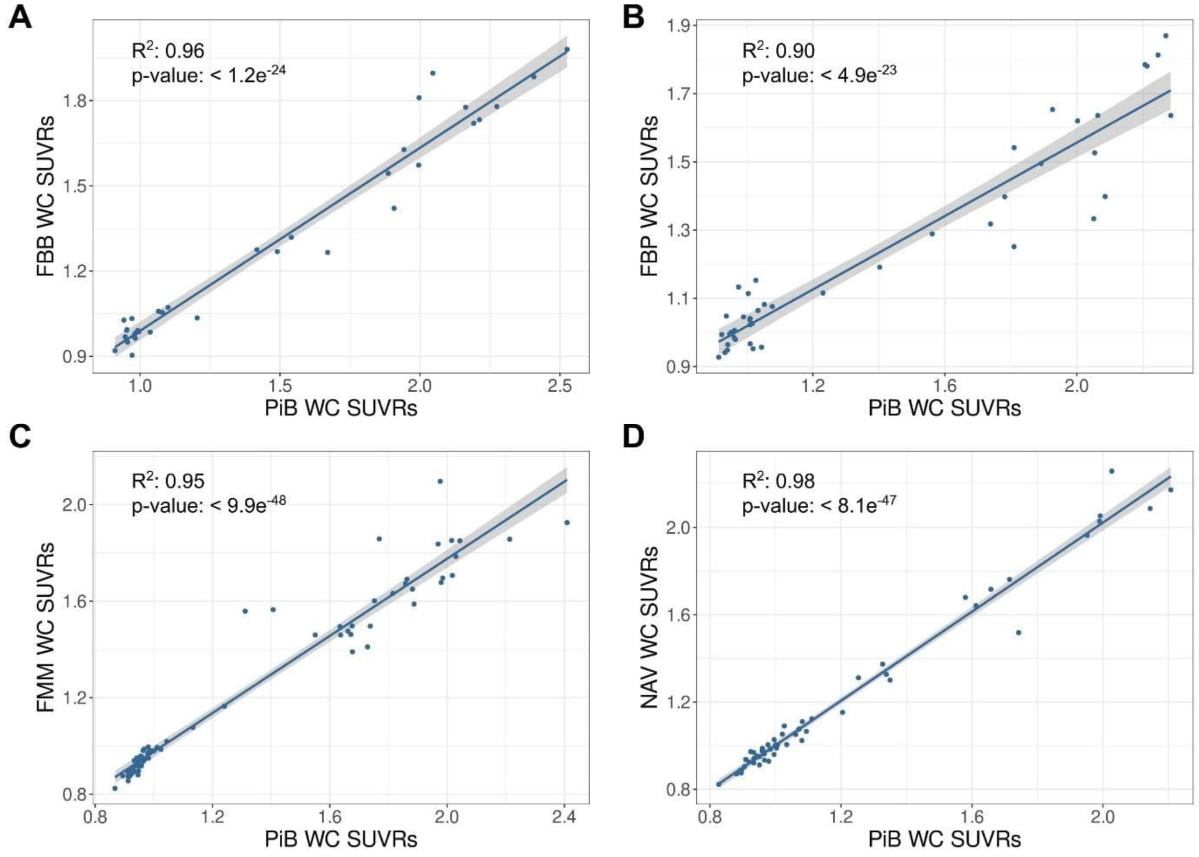


**Figure S2. Head-to-head SUVR correlation for global equation validation.** Scatterplots demonstrating the correlation between paired [11C]PiB SUVRs (x-axes) and the SUVRs of each of the four fluorinated amyloid tracers (y-axes) within the head-to-head validation datasets. Panels correspond to: (A) [18F]Florbetaben (FBB), (B) [18F]Florbetapir (FBP), (C) [18F]Flutemetamol (FMM) and (D) [18F]NAV4694 (NAV). High variance explained across all comparisons ($R^2 \geq 0.90$) confirms that all pre-specified Centiloid Level-2 calibration criteria were successfully met. The blue lines indicate linear fits, and gray shaded areas denote 95% confidence intervals. WC = Whole Cerebellum reference region.

### Figure S3. Regional head-to-head SUVR correlation between [$^{18}$F] tracers and [$^{11}$C]PiB

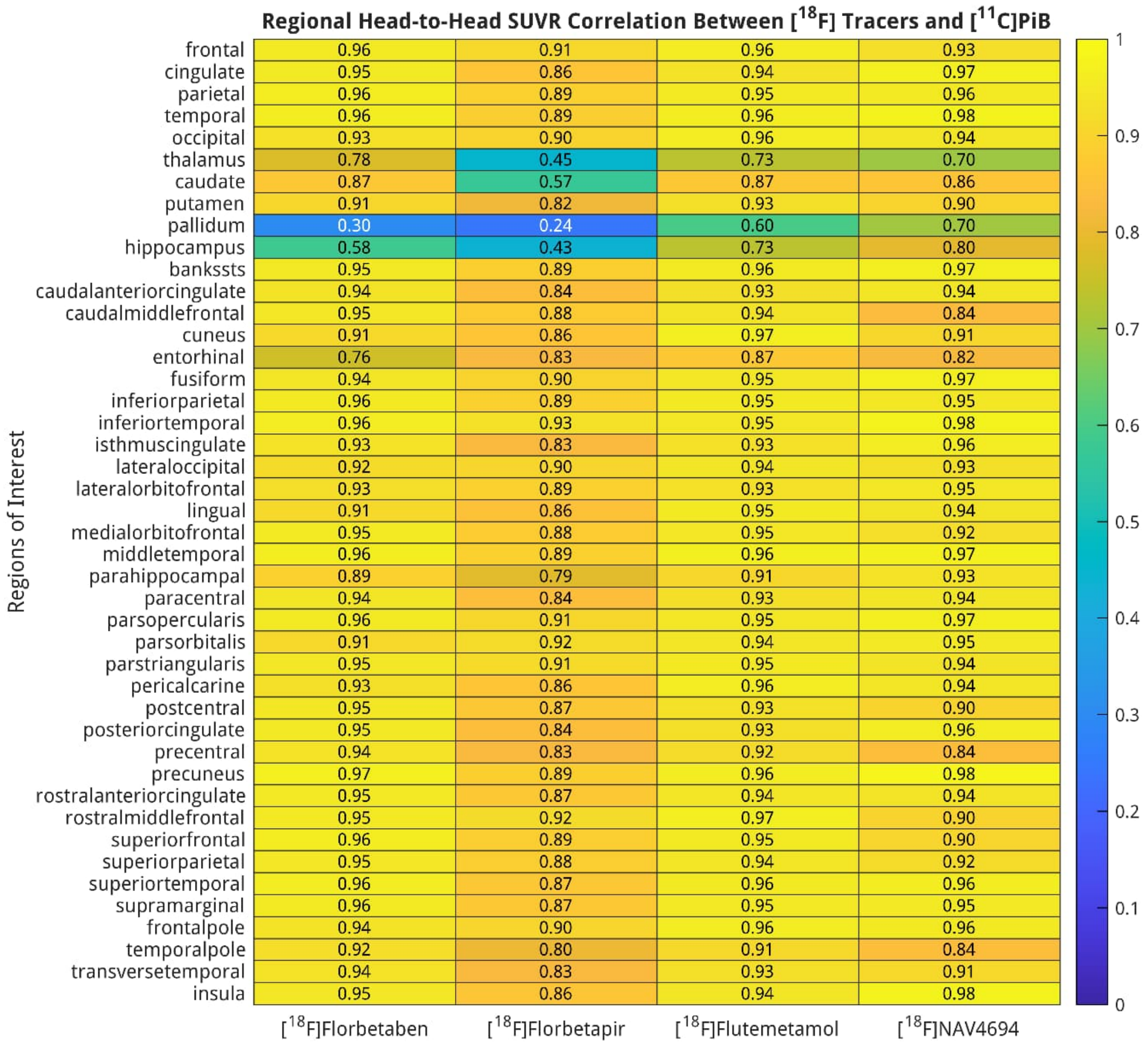


**Figure S3: Regional head-to-head SUVR correlation between [$^{18}$F] tracers and [$^{11}$C]PiB.** Heatmap displaying squared Pearson correlation coefficients ($R^2$) of native Standardized Uptake Value Ratios (SUVR) between each fluorinated tracer ([$^{18}$F]Florbetaben, [$^{18}$F]Florbetapir, [$^{18}$F]Flutemetamol, [$^{18}$F]NAV4694) and the [$^{11}$C]PiB tracer across the different regions of interest from the GAAIN head-to-head datasets. Rows report the ten main target structures (top section: frontal, cingulate, parietal, temporal, occipital, thalamus, caudate, putamen, pallidum, and hippocampus) followed by individual bilateral Desikan-Killiany cortical subregions. Cell values indicate exact regional $R^2$, color-coded according to the scale bar on the right (0 to 1).

**Figure S4. Voxel-wise individual SUVR and Centiloid maps across fluorinated PET radiotracers**

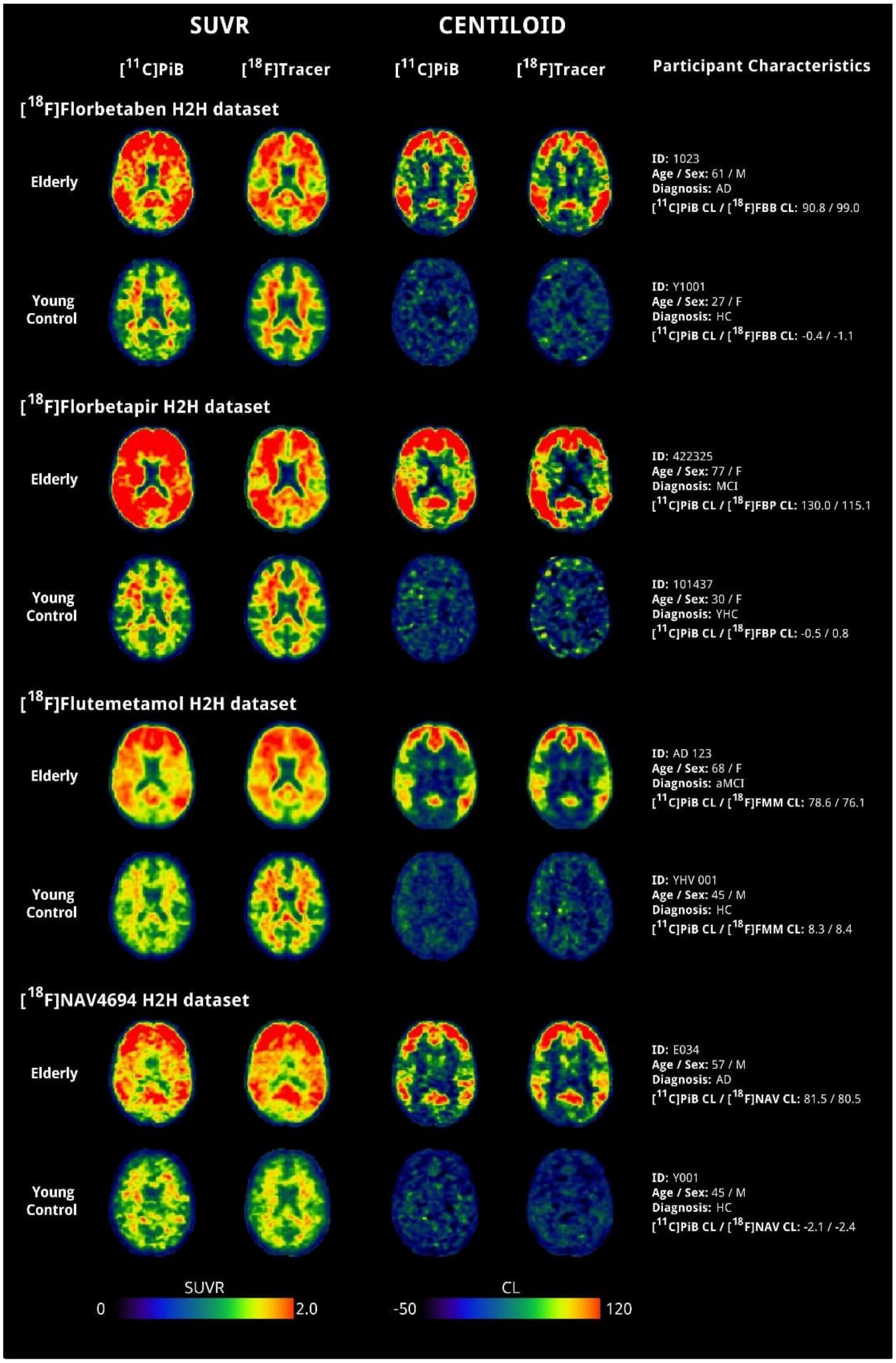


**Figure S4. Individual voxel-wise SUVR and Centiloid maps across fluorinated PET.** Voxel-wise individual parametric maps are displayed on both the native standardized uptake value ratio scale (SUVR, left columns) and the harmonized Centiloid scale (CL, right columns) for one representative elderly participant (top rows) and one amyloid-negative young control (bottom rows) across four head-to-head fluorinated tracer datasets ([$^{18}$F]florbetaben, [$^{18}$F]florbetapir, [$^{18}$F]flutemetamol, and [$^{18}$F]NAV4694). For each tracer dataset, axial slices display the reference [$^{11}$C]PiB acquisition side-by-side with the paired fluorinated acquisition. Centiloid parametric maps (right) were derived using the proposed Level-1 regional zero-anchor recalibration and Level-2 Regional-Intercept calibration models. Color bars represent native SUVR values (0 to 2.0) and standardized Centiloid values (-50 to 120 CL). Additionally, the rightmost column reports participant characteristics: GAAIN ID, age, sex, diagnosis, and official GAAIN-provided global CL values for both [$^{11}$C]PiB and the fluorinated tracer.

### Calibration Parameters

| Tracer | ReferenceRegion | Intercept | Gradient | InterceptCL | GradientCL |
|---|---|---|---|---|---|
| [$^{18}$F]Florbetaben | WC | -0.537 | 1.553 | -163.88 | 167.16 |
| [$^{18}$F]Florbetapir | WC | -0.892 | 1.858 | -202.15 | 200.06 |
| [$^{18}$F]Flutemetamol | WC | -0.221 | 1.251 | -129.94 | 134.68 |
| [$^{18}$F]NAV4694 | WC | 0.019 | 0.979 | -104.07 | 105.45 |
| [$^{11}$C]PiB | WC | 0 | 1 | -106.11 | 107.67 |

**Table S3. Global calibration parameters for cross-tracer SUVR harmonization and Centiloid conversion.** Calibration parameters derived from the standard global cortical target composite using whole cerebellum (WC) as the reference region. The parameters *Gradient* and *Intercept* define linear equations for converting fluorinated radiotracer uptake to [$^{11}$C]PiB-equivalent uptake ($SUVR_{PiB-eq} = Gradient \times SUVR_{F18} + Intercept$). The parameters *GradientCL* and *InterceptCL* provide direct single-step calibration from native fluorinated or $[11C]PiB$ SUVR to global Centiloid values ($CL = GradientCL \times SUVR + InterceptCL$). For [$^{11}$C]PiB, global Centiloid scaling is established directly from the Level-1 young control ($0\ CL$) and Alzheimer disease dementia ($100\ CL$) anchoring cohorts.

**Tables S4 to S8: Regional Centiloid Conversion Equations**

Calibration parameters and linear conversion equations for the Regional-Intercept model across the ten composite regions of interest and the individual Desikan-Killiany atlas regions for each fluorinated radiotracer. Parameters *Gradient* and *Intercept* define linear transformations converting fluorinated radiotracer uptake to [$^{11}$C]PiB-equivalent uptake ($SUVR_{PiB-eq} = Gradient \times SUVR_{F18} + Intercept$). Parameters *GradientCL* and *InterceptCL* enable direct, single-step calibration from native fluorinated SUVR to regional Centiloid values ($CL_r = GradientCL \times SUVR_{F18} + InterceptCL$).

| ROI | Intercept | Gradient | InterceptCL | GradientCL |
|---|---|---|---|---|
| frontal | -0.338 | 1.553 | -162.17 | 167.16 |
| cingulate | -0.337 | 1.553 | -173.91 | 167.16 |
| parietal | -0.362 | 1.553 | -167.98 | 167.16 |
| temporal | -0.339 | 1.553 | -156.94 | 167.16 |
| occipital | -0.384 | 1.553 | -173.25 | 167.16 |
| thalamus | -0.382 | 1.553 | -199.77 | 167.16 |
| caudate | -0.324 | 1.553 | -164.78 | 167.16 |
| putamen | -0.380 | 1.553 | -187.31 | 167.16 |
| pallidum | -0.509 | 1.553 | -238.46 | 167.16 |
| hippocampus | -0.340 | 1.553 | -172.71 | 167.16 |
| ctx-lh-bankssts | -0.427 | 1.553 | -187.52 | 167.16 |
| ctx-lh-caudalanteriorcingulate | -0.350 | 1.553 | -177.17 | 167.16 |
| ctx-lh-caudalmiddlefrontal | -0.370 | 1.553 | -173.39 | 167.16 |
| ctx-lh-cuneus | -0.374 | 1.553 | -176.23 | 167.16 |
| ctx-lh-entorhinal | -0.252 | 1.553 | -145.78 | 167.16 |
| ctx-lh-fusiform | -0.338 | 1.553 | -162.05 | 167.16 |
| ctx-lh-inferiorparietal | -0.379 | 1.553 | -169.12 | 167.16 |
| ctx-lh-inferiortemporal | -0.362 | 1.553 | -158.78 | 167.16 |
| ctx-lh-isthmuscingulate | -0.337 | 1.553 | -177.64 | 167.16 |
| ctx-lh-lateraloccipital | -0.406 | 1.553 | -174.37 | 167.16 |
| ctx-lh-lateralorbitofrontal | -0.365 | 1.553 | -167.81 | 167.16 |
| ctx-lh-lingual | -0.324 | 1.553 | -163.88 | 167.16 |
| ctx-lh-medialorbitofrontal | -0.302 | 1.553 | -156.06 | 167.16 |
| ctx-lh-middletemporal | -0.355 | 1.553 | -157.82 | 167.16 |
| ctx-lh-parahippocampal | -0.288 | 1.553 | -150.10 | 167.16 |
| ctx-lh-paracentral | -0.375 | 1.553 | -178.64 | 167.16 |
| ctx-lh-parsopercularis | -0.345 | 1.553 | -165.78 | 167.16 |
| ctx-lh-parsorbitalis | -0.351 | 1.553 | -153.94 | 167.16 |
| ctx-lh-parstriangularis | -0.357 | 1.553 | -164.84 | 167.16 |
| ctx-lh-pericalcarine | -0.460 | 1.553 | -191.75 | 167.16 |
| ctx-lh-postcentral | -0.359 | 1.553 | -164.39 | 167.16 |
| ctx-lh-posteriorcingulate | -0.347 | 1.553 | -177.88 | 167.16 |

| | | | | |
|---|---|---|---|---|
| ctx-lh-precentral | -0.392 | 1.553 | -176.14 | 167.16 |
| ctx-lh-precuneus | -0.347 | 1.553 | -170.51 | 167.16 |
| ctx-lh-rostralanteriorcingulate | -0.313 | 1.553 | -162.11 | 167.16 |
| ctx-lh-rostralmiddlefrontal | -0.341 | 1.553 | -160.16 | 167.16 |
| ctx-lh-superiorfrontal | -0.327 | 1.553 | -161.60 | 167.16 |
| ctx-lh-superiorparietal | -0.363 | 1.553 | -168.50 | 167.16 |
| ctx-lh-superiortemporal | -0.323 | 1.553 | -156.30 | 167.16 |
| ctx-lh-supramarginal | -0.358 | 1.553 | -164.28 | 167.16 |
| ctx-lh-frontalpole | -0.254 | 1.553 | -128.57 | 167.16 |
| ctx-lh-temporalpole | -0.313 | 1.553 | -146.07 | 167.16 |
| ctx-lh-transversetemporal | -0.330 | 1.553 | -169.38 | 167.16 |
| ctx-lh-insula | -0.327 | 1.553 | -163.86 | 167.16 |
| ctx-rh-bankssts | -0.427 | 1.553 | -187.52 | 167.16 |
| ctx-rh-caudalanteriorcingulate | -0.350 | 1.553 | -177.17 | 167.16 |
| ctx-rh-caudalmiddlefrontal | -0.370 | 1.553 | -173.39 | 167.16 |
| ctx-rh-cuneus | -0.374 | 1.553 | -176.23 | 167.16 |
| ctx-rh-entorhinal | -0.252 | 1.553 | -145.78 | 167.16 |
| ctx-rh-fusiform | -0.338 | 1.553 | -162.05 | 167.16 |
| ctx-rh-inferiorparietal | -0.379 | 1.553 | -169.12 | 167.16 |
| ctx-rh-inferiortemporal | -0.362 | 1.553 | -158.78 | 167.16 |
| ctx-rh-isthmuscingulate | -0.337 | 1.553 | -177.64 | 167.16 |
| ctx-rh-lateraloccipital | -0.406 | 1.553 | -174.37 | 167.16 |
| ctx-rh-lateralorbitofrontal | -0.365 | 1.553 | -167.81 | 167.16 |
| ctx-rh-lingual | -0.324 | 1.553 | -163.88 | 167.16 |
| ctx-rh-medialorbitofrontal | -0.302 | 1.553 | -156.06 | 167.16 |
| ctx-rh-middletemporal | -0.355 | 1.553 | -157.82 | 167.16 |
| ctx-rh-parahippocampal | -0.288 | 1.553 | -150.10 | 167.16 |
| ctx-rh-paracentral | -0.375 | 1.553 | -178.64 | 167.16 |
| ctx-rh-parsopercularis | -0.345 | 1.553 | -165.78 | 167.16 |
| ctx-rh-parsorbitalis | -0.351 | 1.553 | -153.94 | 167.16 |
| ctx-rh-parstriangularis | -0.357 | 1.553 | -164.84 | 167.16 |
| ctx-rh-pericalcarine | -0.460 | 1.553 | -191.75 | 167.16 |
| ctx-rh-postcentral | -0.359 | 1.553 | -164.39 | 167.16 |
| ctx-rh-posteriorcingulate | -0.347 | 1.553 | -177.88 | 167.16 |
| ctx-rh-precentral | -0.392 | 1.553 | -176.14 | 167.16 |
| ctx-rh-precuneus | -0.347 | 1.553 | -170.51 | 167.16 |
| ctx-rh-rostralanteriorcingulate | -0.313 | 1.553 | -162.11 | 167.16 |
| ctx-rh-rostralmiddlefrontal | -0.341 | 1.553 | -160.16 | 167.16 |
| ctx-rh-superiorfrontal | -0.327 | 1.553 | -161.60 | 167.16 |
| ctx-rh-superiorparietal | -0.363 | 1.553 | -168.50 | 167.16 |
| ctx-rh-superiortemporal | -0.323 | 1.553 | -156.30 | 167.16 |

| | | | | |
|---|---|---|---|---|
| ctx-rh-supramarginal | -0.358 | 1.553 | -164.28 | 167.16 |
| ctx-rh-frontalpole | -0.254 | 1.553 | -128.57 | 167.16 |
| ctx-rh-temporalpole | -0.313 | 1.553 | -146.07 | 167.16 |
| ctx-rh-transversetemporal | -0.330 | 1.553 | -169.38 | 167.16 |
| ctx-rh-insula | -0.327 | 1.553 | -163.86 | 167.16 |

**Table S4. Regional calibration parameters for [$^{18}$F]Florbetaben cross-tracer SUVR harmonization and Centiloid conversion.**

| ROI | Intercept | Gradient | InterceptCL | GradientCL |
|---|---|---|---|---|
| frontal | -0.481 | 1.858 | -201.81 | 200.06 |
| cingulate | -0.504 | 1.858 | -218.34 | 200.06 |
| parietal | -0.491 | 1.858 | -205.72 | 200.06 |
| temporal | -0.445 | 1.858 | -189.40 | 200.06 |
| occipital | -0.505 | 1.858 | -210.09 | 200.06 |
| thalamus | -0.521 | 1.858 | -240.02 | 200.06 |
| caudate | -0.463 | 1.858 | -203.33 | 200.06 |
| putamen | -0.564 | 1.858 | -236.66 | 200.06 |
| pallidum | -0.663 | 1.858 | -286.08 | 200.06 |
| hippocampus | -0.468 | 1.858 | -209.53 | 200.06 |
| ctx-lh-bankssts | -0.579 | 1.858 | -231.91 | 200.06 |
| ctx-lh-caudalanteriorcingulate | -0.521 | 1.858 | -223.00 | 200.06 |
| ctx-lh-caudalmiddlefrontal | -0.523 | 1.858 | -216.17 | 200.06 |
| ctx-lh-cuneus | -0.487 | 1.858 | -211.08 | 200.06 |
| ctx-lh-entorhinal | -0.357 | 1.858 | -175.01 | 200.06 |
| ctx-lh-fusiform | -0.493 | 1.858 | -204.34 | 200.06 |
| ctx-lh-inferiorparietal | -0.501 | 1.858 | -206.13 | 200.06 |
| ctx-lh-inferiortemporal | -0.494 | 1.858 | -197.05 | 200.06 |
| ctx-lh-isthmuscingulate | -0.505 | 1.858 | -222.27 | 200.06 |
| ctx-lh-lateraloccipital | -0.515 | 1.858 | -209.52 | 200.06 |
| ctx-lh-lateralorbitofrontal | -0.510 | 1.858 | -208.82 | 200.06 |
| ctx-lh-lingual | -0.469 | 1.858 | -203.51 | 200.06 |
| ctx-lh-medialorbitofrontal | -0.443 | 1.858 | -194.26 | 200.06 |
| ctx-lh-middletemporal | -0.464 | 1.858 | -191.22 | 200.06 |
| ctx-lh-parahippocampal | -0.417 | 1.858 | -185.35 | 200.06 |
| ctx-lh-paracentral | -0.539 | 1.858 | -223.71 | 200.06 |
| ctx-lh-parsopercularis | -0.499 | 1.858 | -208.03 | 200.06 |
| ctx-lh-parsorbitalis | -0.479 | 1.858 | -191.04 | 200.06 |
| ctx-lh-parstriangularis | -0.508 | 1.858 | -206.82 | 200.06 |
| ctx-lh-pericalcarine | -0.586 | 1.858 | -232.06 | 200.06 |
| ctx-lh-postcentral | -0.491 | 1.858 | -202.51 | 200.06 |
| ctx-lh-posteriorcingulate | -0.524 | 1.858 | -224.72 | 200.06 |

| | | | | |
|---|---|---|---|---|
| ctx-lh-precentral | -0.530 | 1.858 | -216.52 | 200.06 |
| ctx-lh-precuneus | -0.495 | 1.858 | -211.51 | 200.06 |
| ctx-lh-rostralanteriorcingulate | -0.466 | 1.858 | -202.98 | 200.06 |
| ctx-lh-rostralmiddlefrontal | -0.480 | 1.858 | -199.22 | 200.06 |
| ctx-lh-superiorfrontal | -0.468 | 1.858 | -200.65 | 200.06 |
| ctx-lh-superiorparietal | -0.478 | 1.858 | -203.51 | 200.06 |
| ctx-lh-superiortemporal | -0.429 | 1.858 | -187.96 | 200.06 |
| ctx-lh-supramarginal | -0.493 | 1.858 | -203.12 | 200.06 |
| ctx-lh-frontalpole | -0.351 | 1.858 | -156.27 | 200.06 |
| ctx-lh-temporalpole | -0.396 | 1.858 | -172.93 | 200.06 |
| ctx-lh-transversetemporal | -0.452 | 1.858 | -204.69 | 200.06 |
| ctx-lh-insula | -0.456 | 1.858 | -200.49 | 200.06 |
| ctx-rh-bankssts | -0.579 | 1.858 | -231.91 | 200.06 |
| ctx-rh-caudalanteriorcingulate | -0.521 | 1.858 | -223.00 | 200.06 |
| ctx-rh-caudalmiddlefrontal | -0.523 | 1.858 | -216.17 | 200.06 |
| ctx-rh-cuneus | -0.487 | 1.858 | -211.08 | 200.06 |
| ctx-rh-entorhinal | -0.357 | 1.858 | -175.01 | 200.06 |
| ctx-rh-fusiform | -0.493 | 1.858 | -204.34 | 200.06 |
| ctx-rh-inferiorparietal | -0.501 | 1.858 | -206.13 | 200.06 |
| ctx-rh-inferiortemporal | -0.494 | 1.858 | -197.05 | 200.06 |
| ctx-rh-isthmuscingulate | -0.505 | 1.858 | -222.27 | 200.06 |
| ctx-rh-lateraloccipital | -0.515 | 1.858 | -209.52 | 200.06 |
| ctx-rh-lateralorbitofrontal | -0.510 | 1.858 | -208.82 | 200.06 |
| ctx-rh-lingual | -0.469 | 1.858 | -203.51 | 200.06 |
| ctx-rh-medialorbitofrontal | -0.443 | 1.858 | -194.26 | 200.06 |
| ctx-rh-middletemporal | -0.464 | 1.858 | -191.22 | 200.06 |
| ctx-rh-parahippocampal | -0.417 | 1.858 | -185.35 | 200.06 |
| ctx-rh-paracentral | -0.539 | 1.858 | -223.71 | 200.06 |
| ctx-rh-parsopercularis | -0.499 | 1.858 | -208.03 | 200.06 |
| ctx-rh-parsorbitalis | -0.479 | 1.858 | -191.04 | 200.06 |
| ctx-rh-parstriangularis | -0.508 | 1.858 | -206.82 | 200.06 |
| ctx-rh-pericalcarine | -0.586 | 1.858 | -232.06 | 200.06 |
| ctx-rh-postcentral | -0.491 | 1.858 | -202.51 | 200.06 |
| ctx-rh-posteriorcingulate | -0.524 | 1.858 | -224.72 | 200.06 |
| ctx-rh-precentral | -0.530 | 1.858 | -216.52 | 200.06 |
| ctx-rh-precuneus | -0.495 | 1.858 | -211.51 | 200.06 |
| ctx-rh-rostralanteriorcingulate | -0.466 | 1.858 | -202.98 | 200.06 |
| ctx-rh-rostralmiddlefrontal | -0.480 | 1.858 | -199.22 | 200.06 |
| ctx-rh-superiorfrontal | -0.468 | 1.858 | -200.65 | 200.06 |
| ctx-rh-superiorparietal | -0.478 | 1.858 | -203.51 | 200.06 |
| ctx-rh-superiortemporal | -0.429 | 1.858 | -187.96 | 200.06 |

| | | | | |
|---|---|---|---|---|
| ctx-rh-supramarginal | -0.493 | 1.858 | -203.12 | 200.06 |
| ctx-rh-frontalpole | -0.351 | 1.858 | -156.27 | 200.06 |
| ctx-rh-temporalpole | -0.396 | 1.858 | -172.93 | 200.06 |
| ctx-rh-transversetemporal | -0.452 | 1.858 | -204.69 | 200.06 |
| ctx-rh-insula | -0.456 | 1.858 | -200.49 | 200.06 |

**Table S5. Regional calibration parameters for [$^{18}$F]Florbetapir cross-tracer SUVR harmonization and Centiloid conversion.**

| ROI | Intercept | Gradient | InterceptCL | GradientCL |
|---|---|---|---|---|
| frontal | -0.167 | 1.251 | -128.14 | 134.67 |
| cingulate | -0.212 | 1.251 | -146.16 | 134.67 |
| parietal | -0.187 | 1.251 | -132.66 | 134.67 |
| temporal | -0.157 | 1.251 | -121.48 | 134.67 |
| occipital | -0.167 | 1.251 | -131.59 | 134.67 |
| thalamus | -0.373 | 1.251 | -186.06 | 134.67 |
| caudate | -0.218 | 1.251 | -139.97 | 134.67 |
| putamen | -0.297 | 1.251 | -163.87 | 134.67 |
| pallidum | -0.516 | 1.251 | -222.89 | 134.67 |
| hippocampus | -0.221 | 1.251 | -145.74 | 134.67 |
| ctx-lh-bankssts | -0.251 | 1.251 | -149.96 | 134.67 |
| ctx-lh-caudalanteriorcingulate | -0.226 | 1.251 | -149.19 | 134.67 |
| ctx-lh-caudalmiddlefrontal | -0.209 | 1.251 | -139.65 | 134.67 |
| ctx-lh-cuneus | -0.164 | 1.251 | -135.82 | 134.67 |
| ctx-lh-entorhinal | -0.116 | 1.251 | -119.22 | 134.67 |
| ctx-lh-fusiform | -0.171 | 1.251 | -128.63 | 134.67 |
| ctx-lh-inferiorparietal | -0.186 | 1.251 | -130.85 | 134.67 |
| ctx-lh-inferiortemporal | -0.167 | 1.251 | -120.80 | 134.67 |
| ctx-lh-isthmuscingulate | -0.220 | 1.251 | -150.94 | 134.67 |
| ctx-lh-lateraloccipital | -0.174 | 1.251 | -129.85 | 134.67 |
| ctx-lh-lateralorbitofrontal | -0.176 | 1.251 | -130.60 | 134.67 |
| ctx-lh-lingual | -0.147 | 1.251 | -129.46 | 134.67 |
| ctx-lh-medialorbitofrontal | -0.166 | 1.251 | -127.96 | 134.67 |
| ctx-lh-middletemporal | -0.150 | 1.251 | -118.57 | 134.67 |
| ctx-lh-parahippocampal | -0.157 | 1.251 | -123.21 | 134.67 |
| ctx-lh-paracentral | -0.220 | 1.251 | -145.62 | 134.67 |
| ctx-lh-parsopercularis | -0.187 | 1.251 | -133.39 | 134.67 |
| ctx-lh-parsorbitalis | -0.124 | 1.251 | -111.94 | 134.67 |
| ctx-lh-parstriangularis | -0.174 | 1.251 | -128.51 | 134.67 |
| ctx-lh-pericalcarine | -0.198 | 1.251 | -141.58 | 134.67 |
| ctx-lh-postcentral | -0.187 | 1.251 | -129.54 | 134.67 |
| ctx-lh-posteriorcingulate | -0.213 | 1.251 | -148.61 | 134.67 |

| | | | | |
|---|---|---|---|---|
| ctx-lh-precentral | -0.227 | 1.251 | -141.16 | 134.67 |
| ctx-lh-precuneus | -0.187 | 1.251 | -137.77 | 134.67 |
| ctx-lh-rostralanteriorcingulate | -0.191 | 1.251 | -135.53 | 134.67 |
| ctx-lh-rostralmiddlefrontal | -0.153 | 1.251 | -123.74 | 134.67 |
| ctx-lh-superiorfrontal | -0.169 | 1.251 | -129.79 | 134.67 |
| ctx-lh-superiorparietal | -0.184 | 1.251 | -132.59 | 134.67 |
| ctx-lh-superiortemporal | -0.164 | 1.251 | -124.38 | 134.67 |
| ctx-lh-supramarginal | -0.194 | 1.251 | -130.61 | 134.67 |
| ctx-lh-frontalpole | -0.075 | 1.251 | -96.22 | 134.67 |
| ctx-lh-temporalpole | -0.135 | 1.251 | -111.93 | 134.67 |
| ctx-lh-transversetemporal | -0.193 | 1.251 | -140.23 | 134.67 |
| ctx-lh-insula | -0.212 | 1.251 | -137.84 | 134.67 |
| ctx-rh-bankssts | -0.251 | 1.251 | -149.96 | 134.67 |
| ctx-rh-caudalanteriorcingulate | -0.226 | 1.251 | -149.19 | 134.67 |
| ctx-rh-caudalmiddlefrontal | -0.209 | 1.251 | -139.65 | 134.67 |
| ctx-rh-cuneus | -0.164 | 1.251 | -135.82 | 134.67 |
| ctx-rh-entorhinal | -0.116 | 1.251 | -119.22 | 134.67 |
| ctx-rh-fusiform | -0.171 | 1.251 | -128.63 | 134.67 |
| ctx-rh-inferiorparietal | -0.186 | 1.251 | -130.85 | 134.67 |
| ctx-rh-inferiortemporal | -0.167 | 1.251 | -120.80 | 134.67 |
| ctx-rh-isthmuscingulate | -0.220 | 1.251 | -150.94 | 134.67 |
| ctx-rh-lateraloccipital | -0.174 | 1.251 | -129.85 | 134.67 |
| ctx-rh-lateralorbitofrontal | -0.176 | 1.251 | -130.60 | 134.67 |
| ctx-rh-lingual | -0.147 | 1.251 | -129.46 | 134.67 |
| ctx-rh-medialorbitofrontal | -0.166 | 1.251 | -127.96 | 134.67 |
| ctx-rh-middletemporal | -0.150 | 1.251 | -118.57 | 134.67 |
| ctx-rh-parahippocampal | -0.157 | 1.251 | -123.21 | 134.67 |
| ctx-rh-paracentral | -0.220 | 1.251 | -145.62 | 134.67 |
| ctx-rh-parsopercularis | -0.187 | 1.251 | -133.39 | 134.67 |
| ctx-rh-parsorbitalis | -0.124 | 1.251 | -111.94 | 134.67 |
| ctx-rh-parstriangularis | -0.174 | 1.251 | -128.51 | 134.67 |
| ctx-rh-pericalcarine | -0.198 | 1.251 | -141.58 | 134.67 |
| ctx-rh-postcentral | -0.187 | 1.251 | -129.54 | 134.67 |
| ctx-rh-posteriorcingulate | -0.213 | 1.251 | -148.61 | 134.67 |
| ctx-rh-precentral | -0.227 | 1.251 | -141.16 | 134.67 |
| ctx-rh-precuneus | -0.187 | 1.251 | -137.77 | 134.67 |
| ctx-rh-rostralanteriorcingulate | -0.191 | 1.251 | -135.53 | 134.67 |
| ctx-rh-rostralmiddlefrontal | -0.153 | 1.251 | -123.74 | 134.67 |
| ctx-rh-superiorfrontal | -0.169 | 1.251 | -129.79 | 134.67 |
| ctx-rh-superiorparietal | -0.184 | 1.251 | -132.59 | 134.67 |
| ctx-rh-superiortemporal | -0.164 | 1.251 | -124.38 | 134.67 |

| | | | | |
|---|---|---|---|---|
| ctx-rh-supramarginal | -0.194 | 1.251 | -130.61 | 134.67 |
| ctx-rh-frontalpole | -0.075 | 1.251 | -96.22 | 134.67 |
| ctx-rh-temporalpole | -0.135 | 1.251 | -111.93 | 134.67 |
| ctx-rh-transversetemporal | -0.193 | 1.251 | -140.23 | 134.67 |
| ctx-rh-insula | -0.212 | 1.251 | -137.84 | 134.67 |

**Table S6. Regional calibration parameters for [$^{18}$F]Flutemetamol cross-tracer SUVR harmonization and Centiloid conversion.**

| ROI | Intercept | Gradient | InterceptCL | GradientCL |
|---|---|---|---|---|
| frontal | 0.026 | 0.979 | -102.84 | 105.45 |
| cingulate | 0.002 | 0.979 | -117.35 | 105.45 |
| parietal | 0.023 | 0.979 | -104.97 | 105.45 |
| temporal | 0.007 | 0.979 | -99.62 | 105.45 |
| occipital | 0.007 | 0.979 | -108.34 | 105.45 |
| thalamus | 0.044 | 0.979 | -131.26 | 105.45 |
| caudate | -0.006 | 0.979 | -111.25 | 105.45 |
| putamen | 0.006 | 0.979 | -123.21 | 105.45 |
| pallidum | 0.003 | 0.979 | -153.10 | 105.45 |
| hippocampus | -0.046 | 0.979 | -120.78 | 105.45 |
| ctx-lh-bankssts | 0.014 | 0.979 | -114.69 | 105.45 |
| ctx-lh-caudalanteriorcingulate | 0.000 | 0.979 | -118.73 | 105.45 |
| ctx-lh-caudalmiddlefrontal | 0.065 | 0.979 | -104.65 | 105.45 |
| ctx-lh-cuneus | 0.024 | 0.979 | -111.22 | 105.45 |
| ctx-lh-entorhinal | -0.085 | 0.979 | -112.60 | 105.45 |
| ctx-lh-fusiform | -0.029 | 0.979 | -108.65 | 105.45 |
| ctx-lh-inferiorparietal | 0.019 | 0.979 | -103.78 | 105.45 |
| ctx-lh-inferiortemporal | -0.034 | 0.979 | -101.86 | 105.45 |
| ctx-lh-isthmuscingulate | -0.015 | 0.979 | -122.87 | 105.45 |
| ctx-lh-lateraloccipital | -0.003 | 0.979 | -106.77 | 105.45 |
| ctx-lh-lateralorbitofrontal | -0.040 | 0.979 | -111.13 | 105.45 |
| ctx-lh-lingual | 0.011 | 0.979 | -108.55 | 105.45 |
| ctx-lh-medialorbitofrontal | 0.001 | 0.979 | -105.52 | 105.45 |
| ctx-lh-middletemporal | -0.003 | 0.979 | -98.77 | 105.45 |
| ctx-lh-parahippocampal | -0.034 | 0.979 | -105.61 | 105.45 |
| ctx-lh-paracentral | 0.022 | 0.979 | -113.63 | 105.45 |
| ctx-lh-parsopercularis | 0.025 | 0.979 | -105.50 | 105.45 |
| ctx-lh-parsorbitalis | -0.070 | 0.979 | -102.62 | 105.45 |
| ctx-lh-parstriangularis | 0.002 | 0.979 | -104.93 | 105.45 |
| ctx-lh-pericalcarine | 0.027 | 0.979 | -112.03 | 105.45 |
| ctx-lh-postcentral | 0.032 | 0.979 | -100.96 | 105.45 |
| ctx-lh-posteriorcingulate | 0.019 | 0.979 | -117.84 | 105.45 |

| | | | | |
|---|---|---|---|---|
| ctx-lh-precentral | 0.047 | 0.979 | -105.62 | 105.45 |
| ctx-lh-precuneus | 0.020 | 0.979 | -110.46 | 105.45 |
| ctx-lh-rostralanteriorcingulate | 0.005 | 0.979 | -109.27 | 105.45 |
| ctx-lh-rostralmiddlefrontal | 0.041 | 0.979 | -98.83 | 105.45 |
| ctx-lh-superiorfrontal | 0.055 | 0.979 | -101.24 | 105.45 |
| ctx-lh-superiorparietal | 0.040 | 0.979 | -103.66 | 105.45 |
| ctx-lh-superiortemporal | 0.016 | 0.979 | -100.57 | 105.45 |
| ctx-lh-supramarginal | 0.012 | 0.979 | -103.20 | 105.45 |
| ctx-lh-frontalpole | 0.005 | 0.979 | -85.63 | 105.45 |
| ctx-lh-temporalpole | -0.078 | 0.979 | -102.02 | 105.45 |
| ctx-lh-transversetemporal | 0.045 | 0.979 | -109.45 | 105.45 |
| ctx-lh-insula | 0.007 | 0.979 | -108.47 | 105.45 |
| ctx-rh-bankssts | 0.014 | 0.979 | -114.69 | 105.45 |
| ctx-rh-caudalanteriorcingulate | 0.000 | 0.979 | -118.73 | 105.45 |
| ctx-rh-caudalmiddlefrontal | 0.065 | 0.979 | -104.65 | 105.45 |
| ctx-rh-cuneus | 0.024 | 0.979 | -111.22 | 105.45 |
| ctx-rh-entorhinal | -0.085 | 0.979 | -112.60 | 105.45 |
| ctx-rh-fusiform | -0.029 | 0.979 | -108.65 | 105.45 |
| ctx-rh-inferiorparietal | 0.019 | 0.979 | -103.78 | 105.45 |
| ctx-rh-inferiortemporal | -0.034 | 0.979 | -101.86 | 105.45 |
| ctx-rh-isthmuscingulate | -0.015 | 0.979 | -122.87 | 105.45 |
| ctx-rh-lateraloccipital | -0.003 | 0.979 | -106.77 | 105.45 |
| ctx-rh-lateralorbitofrontal | -0.040 | 0.979 | -111.13 | 105.45 |
| ctx-rh-lingual | 0.011 | 0.979 | -108.55 | 105.45 |
| ctx-rh-medialorbitofrontal | 0.001 | 0.979 | -105.52 | 105.45 |
| ctx-rh-middletemporal | -0.003 | 0.979 | -98.77 | 105.45 |
| ctx-rh-parahippocampal | -0.034 | 0.979 | -105.61 | 105.45 |
| ctx-rh-paracentral | 0.022 | 0.979 | -113.63 | 105.45 |
| ctx-rh-parsopercularis | 0.025 | 0.979 | -105.50 | 105.45 |
| ctx-rh-parsorbitalis | -0.070 | 0.979 | -102.62 | 105.45 |
| ctx-rh-parstriangularis | 0.002 | 0.979 | -104.93 | 105.45 |
| ctx-rh-pericalcarine | 0.027 | 0.979 | -112.03 | 105.45 |
| ctx-rh-postcentral | 0.032 | 0.979 | -100.96 | 105.45 |
| ctx-rh-posteriorcingulate | 0.019 | 0.979 | -117.84 | 105.45 |
| ctx-rh-precentral | 0.047 | 0.979 | -105.62 | 105.45 |
| ctx-rh-precuneus | 0.020 | 0.979 | -110.46 | 105.45 |
| ctx-rh-rostralanteriorcingulate | 0.005 | 0.979 | -109.27 | 105.45 |
| ctx-rh-rostralmiddlefrontal | 0.041 | 0.979 | -98.83 | 105.45 |
| ctx-rh-superiorfrontal | 0.055 | 0.979 | -101.24 | 105.45 |
| ctx-rh-superiorparietal | 0.040 | 0.979 | -103.66 | 105.45 |
| ctx-rh-superiortemporal | 0.016 | 0.979 | -100.57 | 105.45 |

| | | | | |
|---|---|---|---|---|
| ctx-rh-supramarginal | 0.012 | 0.979 | -103.20 | 105.45 |
| ctx-rh-frontalpole | 0.005 | 0.979 | -85.63 | 105.45 |
| ctx-rh-temporalpole | -0.078 | 0.979 | -102.02 | 105.45 |
| ctx-rh-transversetemporal | 0.045 | 0.979 | -109.45 | 105.45 |
| ctx-rh-insula | 0.007 | 0.979 | -108.47 | 105.45 |

**Table S7. Regional calibration parameters for [$^{18}$F]NAV4694 cross-tracer SUVR harmonization and Centiloid conversion.**

| ROI | InterceptCL | GradientCL |
|---|---|---|
| frontal | -105.60 | 107.67 |
| cingulate | -117.60 | 107.67 |
| parietal | -107.43 | 107.67 |
| temporal | -100.32 | 107.67 |
| occipital | -109.13 | 107.67 |
| thalamus | -135.87 | 107.67 |
| caudate | -110.67 | 107.67 |
| putamen | -123.86 | 107.67 |
| pallidum | -153.43 | 107.67 |
| hippocampus | -115.95 | 107.67 |
| ctx-lh-bankssts | -116.17 | 107.67 |
| ctx-lh-caudalanteriorcingulate | -118.72 | 107.67 |
| ctx-lh-caudalmiddlefrontal | -111.55 | 107.67 |
| ctx-lh-cuneus | -113.74 | 107.67 |
| ctx-lh-entorhinal | -103.61 | 107.67 |
| ctx-lh-fusiform | -105.62 | 107.67 |
| ctx-lh-inferiorparietal | -105.81 | 107.67 |
| ctx-lh-inferiortemporal | -98.32 | 107.67 |
| ctx-lh-isthmuscingulate | -121.25 | 107.67 |
| ctx-lh-lateraloccipital | -106.47 | 107.67 |
| ctx-lh-lateralorbitofrontal | -106.86 | 107.67 |
| ctx-lh-lingual | -109.72 | 107.67 |
| ctx-lh-medialorbitofrontal | -105.66 | 107.67 |
| ctx-lh-middletemporal | -98.42 | 107.67 |
| ctx-lh-parahippocampal | -102.01 | 107.67 |
| ctx-lh-paracentral | -115.93 | 107.67 |
| ctx-lh-parsopercularis | -108.18 | 107.67 |
| ctx-lh-parsorbitalis | -95.24 | 107.67 |
| ctx-lh-parstriangularis | -105.14 | 107.67 |
| ctx-lh-pericalcarine | -114.90 | 107.67 |
| ctx-lh-postcentral | -104.30 | 107.67 |
| ctx-lh-posteriorcingulate | -119.86 | 107.67 |

| | | |
|---|---|---|
| ctx-lh-precentral | -110.55 | 107.67 |
| ctx-lh-precuneus | -112.52 | 107.67 |
| ctx-lh-rostralanteriorcingulate | -109.74 | 107.67 |
| ctx-lh-rostralmiddlefrontal | -103.18 | 107.67 |
| ctx-lh-superiorfrontal | -107.02 | 107.67 |
| ctx-lh-superiorparietal | -107.86 | 107.67 |
| ctx-lh-superiortemporal | -102.23 | 107.67 |
| ctx-lh-supramarginal | -104.47 | 107.67 |
| ctx-lh-frontalpole | -86.12 | 107.67 |
| ctx-lh-temporalpole | -93.78 | 107.67 |
| ctx-lh-transversetemporal | -114.21 | 107.67 |
| ctx-lh-insula | -109.25 | 107.67 |
| ctx-rh-bankssts | -116.17 | 107.67 |
| ctx-rh-caudalanteriorcingulate | -118.72 | 107.67 |
| ctx-rh-caudalmiddlefrontal | -111.55 | 107.67 |
| ctx-rh-cuneus | -113.74 | 107.67 |
| ctx-rh-entorhinal | -103.61 | 107.67 |
| ctx-rh-fusiform | -105.62 | 107.67 |
| ctx-rh-inferiorparietal | -105.81 | 107.67 |
| ctx-rh-inferiortemporal | -98.32 | 107.67 |
| ctx-rh-isthmuscingulate | -121.25 | 107.67 |
| ctx-rh-lateraloccipital | -106.47 | 107.67 |
| ctx-rh-lateralorbitofrontal | -106.86 | 107.67 |
| ctx-rh-lingual | -109.72 | 107.67 |
| ctx-rh-medialorbitofrontal | -105.66 | 107.67 |
| ctx-rh-middletemporal | -98.42 | 107.67 |
| ctx-rh-parahippocampal | -102.01 | 107.67 |
| ctx-rh-paracentral | -115.93 | 107.67 |
| ctx-rh-parsopercularis | -108.18 | 107.67 |
| ctx-rh-parsorbitalis | -95.24 | 107.67 |
| ctx-rh-parstriangularis | -105.14 | 107.67 |
| ctx-rh-pericalcarine | -114.90 | 107.67 |
| ctx-rh-postcentral | -104.30 | 107.67 |
| ctx-rh-posteriorcingulate | -119.86 | 107.67 |
| ctx-rh-precentral | -110.55 | 107.67 |
| ctx-rh-precuneus | -112.52 | 107.67 |
| ctx-rh-rostralanteriorcingulate | -109.74 | 107.67 |
| ctx-rh-rostralmiddlefrontal | -103.18 | 107.67 |
| ctx-rh-superiorfrontal | -107.02 | 107.67 |
| ctx-rh-superiorparietal | -107.86 | 107.67 |
| ctx-rh-superiortemporal | -102.23 | 107.67 |

| | | |
|---|---|---|
| ctx-rh-supramarginal | -104.47 | 107.67 |
| ctx-rh-frontalpole | -86.12 | 107.67 |
| ctx-rh-temporalpole | -93.78 | 107.67 |
| ctx-rh-transversetemporal | -114.21 | 107.67 |
| ctx-rh-insula | -109.25 | 107.67 |

**Table S8. Regional calibration parameters for [$^{11}$C]PiB SUVR to Centiloid conversion.**

### Code and Computational Resources

To facilitate reproducibility and implementation of the regional and voxel-wise Centiloid framework, complete computational code and supporting files are made publicly available at:

https://github.com/Erica-Silvestri/AmyloidPET-Regional-Centiloid-Conversion.git

Available resources include:

1. Regional Conversion Scripts: Python implementation for applying the Regional-Intercept calibration equations to regional SUVR measurements, generating regional Centiloid values for the ten primary ROIs and individual Desikan-Killiany atlas regions.
2. Voxel-Wise Conversion Scripts: Python-based pipelines for voxel-wise parametric mapping, including:
    - Implementation of Level-1 regional zero-anchor recalibration at the voxel level
    - Application of Level-2 Regional-Intercept calibration equations
    - Generation of parametric Centiloid maps in standard MNI152 space
3. Standard Space Reference Maps: Pre-computed MNI152 2 $mm^3$ resolution maps of:
    - Level-1 regional intercept values for each tracer
    - Global slope values for cross-tracer conversion
    - These maps enable direct voxel-wise conversion without requiring regional equation re-estimation
4. Calibration Parameter Files: Complete tables of regional slopes, intercepts, and conversion equations for all four fluorinated tracers ([18F]florbetaben, [$^{18}$F]florbetapir, [$^{18}$F]flutemetamol, [$^{18}$F]NAV4694) in .csv format.
